# Mapping Twenty Years of Antimicrobial Resistance Research Trends

**DOI:** 10.1101/2021.03.01.433375

**Authors:** C.F. Luz, J.M. van Niekerk, J. Keizer, N. Beerlage-de Jong, L.M.A. Braakman-Jansen, A. Stein, B. Sinha, J.E.W.C. van Gemert-Pijnen, C. Glasner

**Author notes:** Correspondence to: Christian Luz, University of Groningen, University Medical Center Groningen, Department of Medical Microbiology and Infection Prevention, Hanzeplein 1 EB 80, 9713GZ Groningen, The Netherlands. Equal contribution (C.F. Luz, J.M. van Niekerk).

## Abstract

**Background:** Antimicrobial resistance (AMR) is a global threat to health and healthcare. In response to the growing AMR burden, research funding also increased. However, a comprehensive overview of the research output, including conceptual, temporal, and geographical trends, is missing. Therefore, this study uses topic modelling, a machine learning approach, to reveal the scientific evolution of AMR research and its trends, and provides an interactive user interface for further analyses.

**Methods:** Structural topic modelling (STM) was applied on a text corpus resulting from a PubMed query comprising AMR articles (1999-2018). A topic network was established and topic trends were analysed by frequency, proportion, and importance over time and space.

**Findings:** In total, 88 topics were identified in 158616 articles from 166 countries. AMR publications increased by 450% between 1999 and 2018, emphasizing the vibrancy of the field. Prominent topics in 2018 were *Strategies for emerging resistances and diseases, Nanoparticles*, and *Stewardship*. Emerging topics included *Water and environment*, and *Sequencing*. Geographical trends showed prominence of *Multidrug-resistant tuberculosis (MDR-TB)* in the WHO African Region, corresponding with the MDR-TB burden. China and India were growing contributors in recent years, following the United States of America as overall lead contributor.

**Interpretation:** This study provides a comprehensive overview of the AMR research output thereby revealing the AMR research response to the increased AMR burden. Both the results and the publicly available interactive database serve as a base to inform and optimise future research.

**Funding:** INTERREG-VA EurHealth-1Health (202085); European Commission Horizon 2020 Framework

**Research in context:** *Evidence before this study:* Prior to this study, PubMed, Web of Science, Scopus, and IEEE Xplore were queried to find studies providing a conceptual overview of antimicrobial resistance (AMR) research over time and space. The search string included keywords (“antimicrobial” OR antibiotic*) AND (resistan*) AND (“science mapping” OR bibliometric OR scientometric) in the title and abstract and focused on articles published before 2019 without language restrictions. Few studies were found relying on scientometric and bibliometric methods to assess either subfields of AMR research (e.g., AMR among uropathogens) or AMR-related fields (e.g., microbiology). No studies were found that focus on the entire AMR field. Therefore, this science mapping study using topic modelling was performed to provide an overview of the AMR field by identifying and assessing topics, trends, and geographical differences over time.

*Added value of this study:* To the best of our knowledge, this study is the first to use a science mapping approach to provide a comprehensive overview of the entire AMR research field, covering over 150 thousand articles published between 1999 and 2018. Our findings revealed important (e.g., *Strategies for emerging resistances and diseases, Nanoparticles*, and *Stewardship*) and emerging (e.g., *Water and environment*, and *Sequencing*) topics in AMR research. Lastly, this study resulted in an interactive user interface where all data are presented for further analyses.

*Implications of all the available evidence:* Our comprehensive overview of the AMR field, including its conceptual structure, and temporal and geographical trends revealed the response of the research community to the AMR burden. The results and the openly available supporting data provide the base to guide future funding and research directions to tackle AMR.

## Introduction

Antimicrobial resistance (AMR) is challenging health and healthcare globally. The burden gradually increased over time and recent reports depict extreme predictions, although global estimates are difficult to derive.^1,2^ Previously well-treatable infections require new therapeutic strategies, while already difficult-to-treat diseases have developed extensive resistance, e.g., multidrug-resistant tuberculosis (MDR-TB). International policy bodies and governments have put AMR high on the political agenda and call for more research to ease the AMR rise.^3,4^ Hence, various dedicated research funds have been allocated across countries.^5–7^ While the global AMR burden and research funding are increasing, the response in AMR research output (i.e., scientific evolution) remains unknown; a holistic view on the entire field and its structure is lacking.

Neither the AMR burden nor the appropriate financial resources are equally distributed.^1,2^ Thus, providing a comprehensive picture requires assessing geographical differences. The conceptual structure of AMR research is highly heterogeneous due to its cross-disciplinary nature, making it difficult to grasp the overall picture and interrelatedness of research topics.^8^ However, the structure of the AMR field is essential to identify temporal and geographical trends, assess funding effects, and help guide future research and funding.

Identifying research topics and trends within an entire field is challenging. The amount of publications can hardly be overseen by single individuals anymore. Few studies addressed trends in AMR-related research based on scientometric and bibliometric approaches (i.e., by quantitative means). They focused on the global research output on AMR among uropathogens, carbapenem resistance, AMR in the environment, and AMR history.^9–12^ Other studies aimed at broader levels by identifying research topics in microbiology between 1945-2016 and 2012-2016.^13,14^ These approaches were either too narrow by investigating merely parts of the AMR field or too broad to identify AMR and related subgroups. So far, no comprehensive study is available to provide information on global AMR research activities.

Data-driven, computational approaches can provide solutions to the challenge of identifying topics and trends in texts. Particularly, topic modelling has been used to study entire research fields before.^15^ Topic modelling is a statistical approach, which enables semi-automated topic discovery and exploration within texts.^16,17^ The underlying assumption is that a text consists of one or several topics made up of various words which are likely to co-occur and define the respective topic. Today’s available computational resources allow for applying this unsupervised machine learning technique to large collections of documents.

Studying the entire body of scientific literature on AMR in bacteria and fungi is unprecedented. Our study uses topic modelling to provide an overview of the AMR field by identifying and assessing topics, trends, and geographical differences over time. The study also lays the groundwork for further analyses by providing an interactive user interface, which provides guidance for future research directions in AMR.

## Methods

### Data & Search String

Data was retrieved from the PubMed database using a comprehensive search string to reflect the entire AMR field accessed through the PubMed API.^18–20^ The search string was: (“Anti-Bacterial Agents”[Mesh] OR Anti-Bacterial*[tiab] OR antibacterial*[tiab] OR antibiotic*[tiab] OR antimicrobial*[tiab] OR antimycobacterial*[tiab] OR “Antifungal Agents”[Mesh] OR Antifungal*[tiab] or anti-fungal*[tiab]) AND (“Drug Resistance”[Mesh] OR resistan*[tiab] OR “Microbial Sensitivity Tests”[Mesh]). The search results formed the text corpus of this study.

The extracted variables were: PMID number, author names, affiliations, title, abstract, journal, and database entry year. Records were included if they were published between 1999 and 2018, included an English abstract, and were not part of a set of exclusion criteria for article types (Figure 1). Records’ citations were downloaded from the NCBI Entrez database.

**Figure 1.**
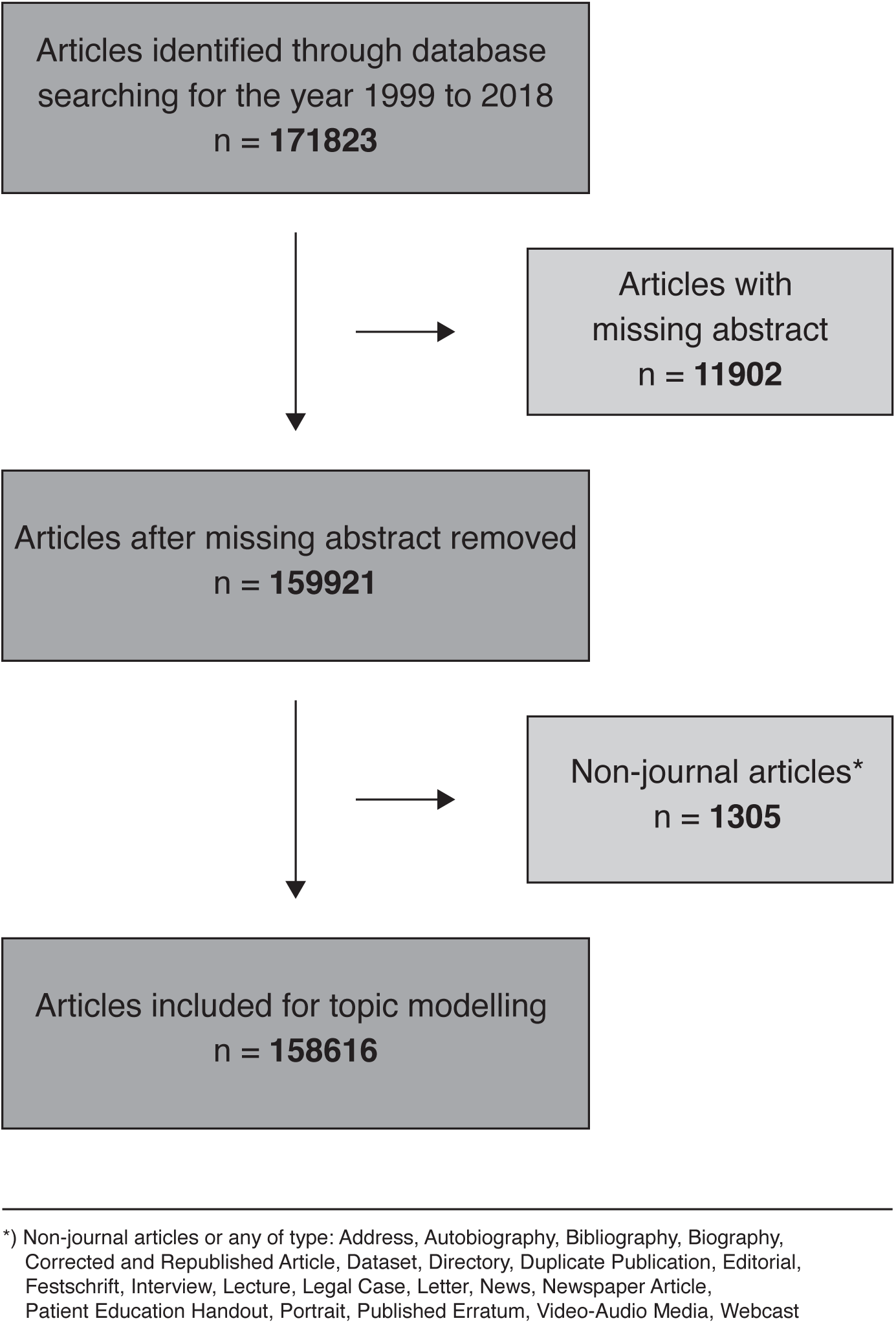
PRISMA flow diagram of included and excluded articles.

### Data pre-processing

Title and abstract were merged into one text variable, excluding non-word and character strings with less than two characters. Generic and domain-specific (Appendix S1) stopwords were removed from the text.^21^ Finally, all text was stemmed using the snowball stemming algorithm.^21^ First author countries were extracted from the affiliation data and complementary grouped into World Health Organization (WHO) regions.

Centrality measures can be used to quantify the extent to which articles influenced each other using network structures^22^. This study considered four measures of centrality (Degree, H-index, PageRank, and Eigenvector) to assess directional relationships of citations between the articles. The PageRank measure was used to determine the importance of countries and topics over time as it had the highest contribution according to principal component analysis.^23^

### Topic modelling

We used structural topic modelling (STM), which extends classical topic modelling (details in Appendix S2) and is available through the stm R package.^20,24^ STM modelling has previously been applied to identify topics in scientific literature.^15–17^ STM modelling is generative similar to latent Dirichlet allocation (LDA) but with the added benefit of including document-level covariates. We used publication year, citation count, PageRank, and first author country as document-level covariates. The output of a topic model is a collection of *bag-of-words* where each bag consists of words that constitute a topic.^25^ We set a broad initial range for the number of bags/topics (*K*; range 15-205) and chose the optimal number by considering how well each number of topics represents the text corpus both quantitatively and qualitatively. The optimal number for *K* was identified by considering the semantic coherence and exclusivity per model of *K* topics and assessing the topics’ interpretabilit.^15–17^

Each model’s output in the identified range of *K* was assessed. Two researchers (CFL, CG) independently assessed all topics based on the associated terms and generated a describing name per topic. Topic names were further refined by scanning titles and abstracts of five highly associated articles per topic and five important articles per topic by PageRank. No topic name was assigned if this process did not converge to a meaningful name. Consensus was reached when both researchers differed in their generated topic names. Five independent AMR researchers reviewed this process and verified the generated topic names. The final model was chosen based on the highest number of topics with an assigned topic name. Each article was assigned the topic name of the topic comprising the highest proportion of the article’s text. Topics in the final model were inductively coded into thematic groups to navigate the results.

### Topic investigation

Topics were assessed with two different objectives: 1) analysing topic relationships; 2) identifying trends by frequency, proportion, and PageRank (importance) over time and space. Topic relationships were assessed using two data sources: 1) within-text-corpus citations for the included publications; 2) topic co-occurrence per article. These data were clustered using 1) hierarchical clustering with Ward’s minimum variance method and 2) topic correlation estimation.^26^ Publication bursts, i.e., the least amount of years comprising 50% of all publications starting at the earliest year possible, were calculated per topic.

### Data availability and interactive user interface

This study generated substantial amounts of data that enable detailed analyses. The results in this manuscript present only selected highlights from these data. To repeat this study’s analyses and to enable further analyses, an interactive web-based application was developed (https://topicsinamr.shinyapps.io/amr_topics/). Additionally, individual articles can be searched and assessed and the topic model can be leveraged to evaluate texts from new articles not included in this study. Moreover, the data used and generated in this study is openly available under (https://osf.io/j3d65/). All analyses and the application development were performed in R.^20^

## Results

In total, 158616 articles were included, showing a steady increase over the past 20 years (8·5% nominal annual increase) (Figure 2). In 2018, 14547 articles were published, an increase of 450% compared to 1999. The topic modelling process using the optimal number of *K*= 95 topics resulted in 88 named topics, covering 152780 articles (96·3%). All topics sorted by thematic groups are presented in Figure 3, including trend lines in publication frequency, annual proportion, and PageRank. The cluster analysis based on topic co-occurrence correlation revealed a tightly connected network of the AMR field (Figure 4).

**Figure 2.**
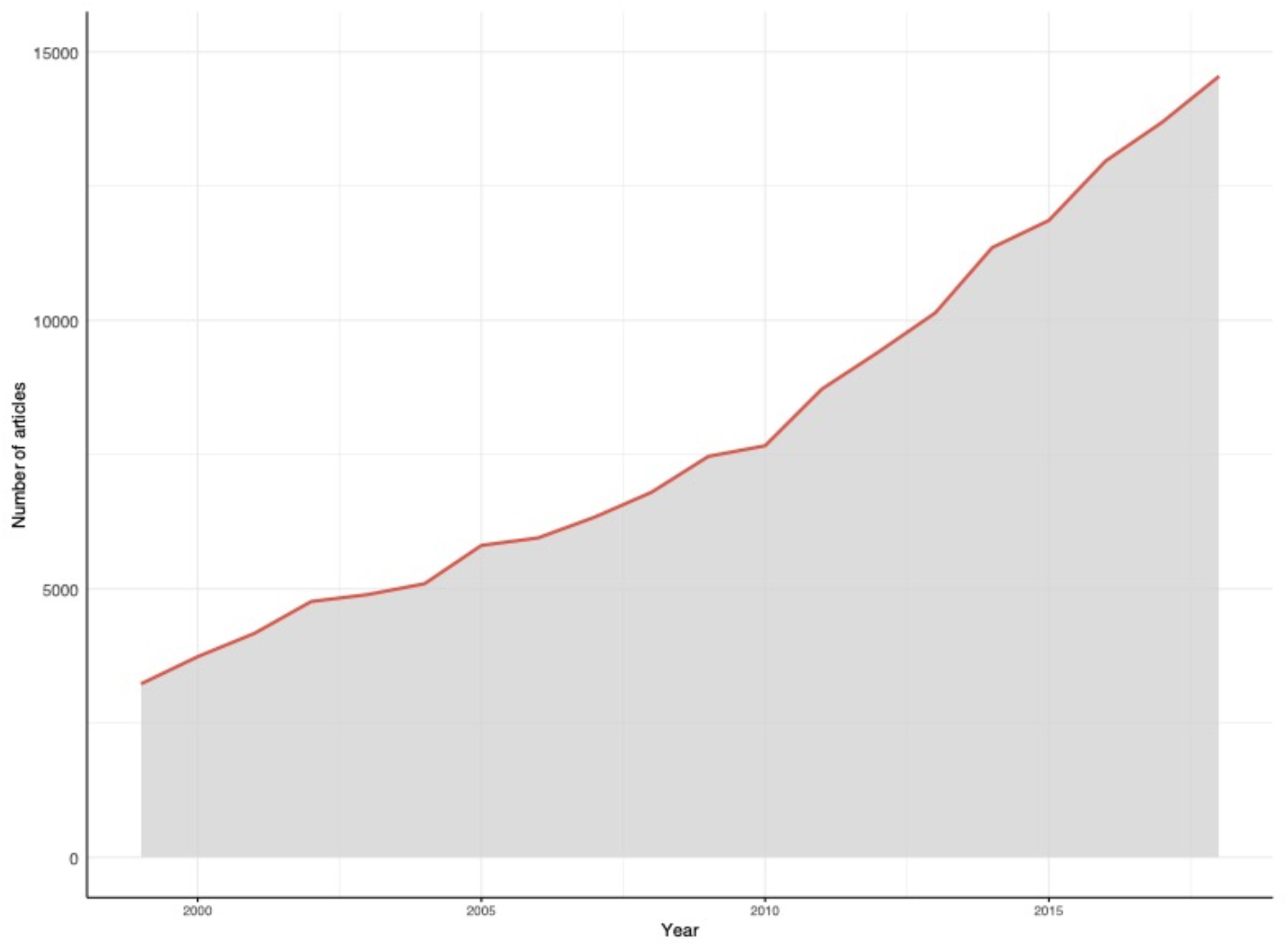
Total number of antimicrobial resistance (AMR) related publications indexed on PubMed per year (1999-2018) based on the applied search string: (“Anti-Bacterial Agents” [Mesh] OR Anti-Bacterial* [tiab] OR antibacterial* [tiab] OR antibiotic* [tiab] OR antimicrobial* [tiab] OR antimycobacterial* [tiab] OR “Antifungal Agents”[Mesh] OR Antifungal* [tiab] or anti-fungal* [tiab]) AND (“Drug Resistance”[Mesh] OR resistan* [tiab] OR “Microbial Sensitivity Tests”[Mesh]).

**Figure 3.**
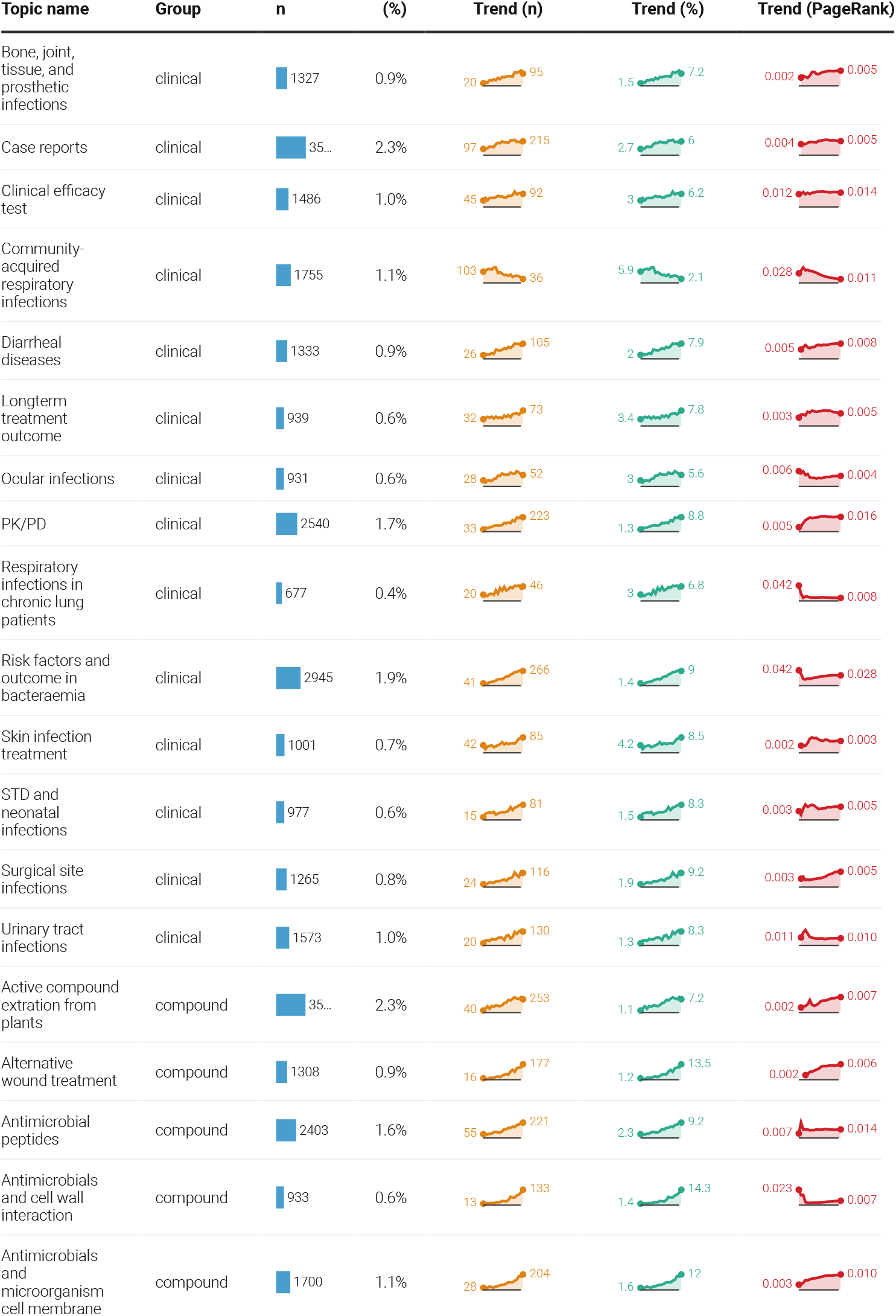
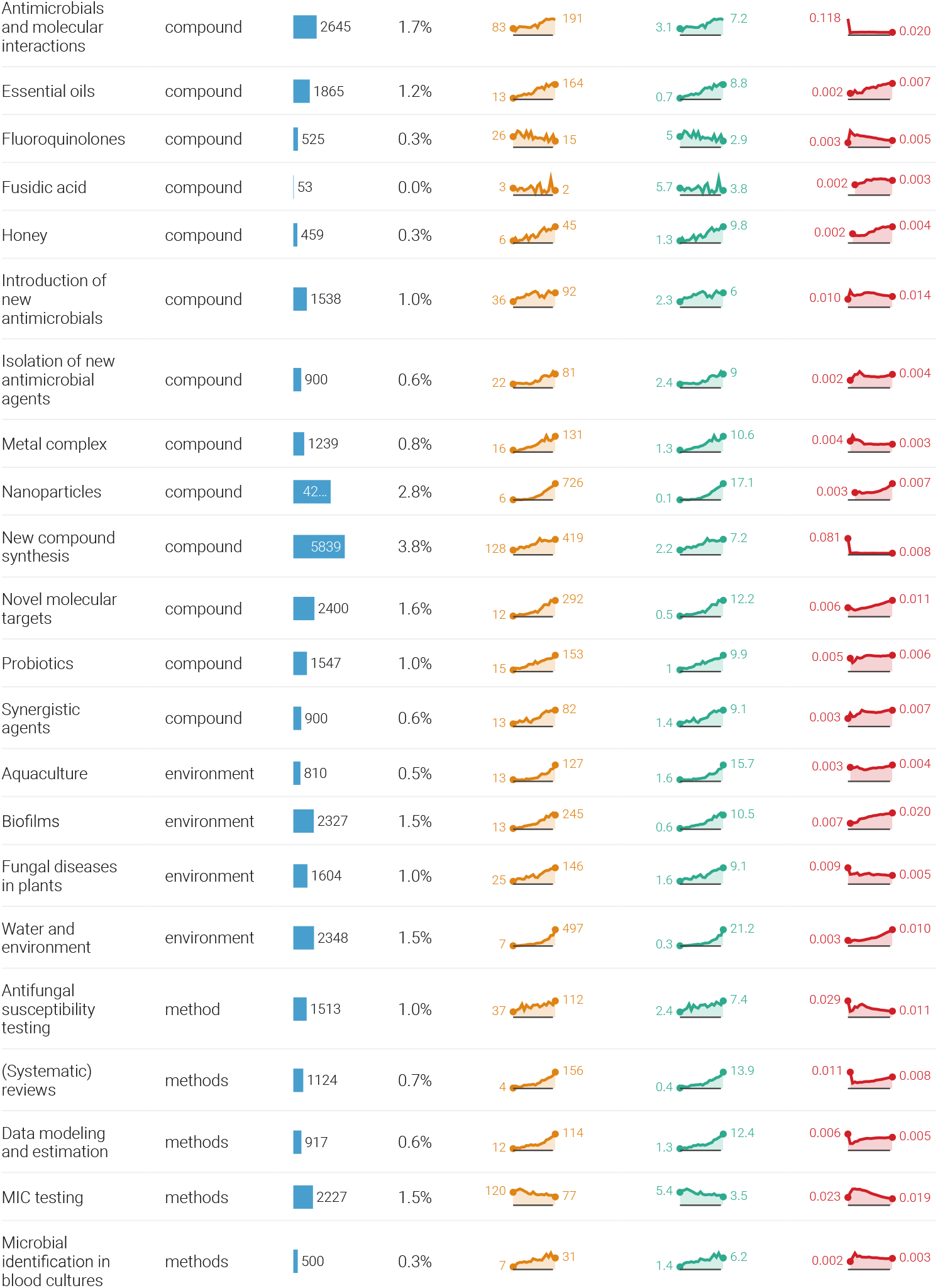
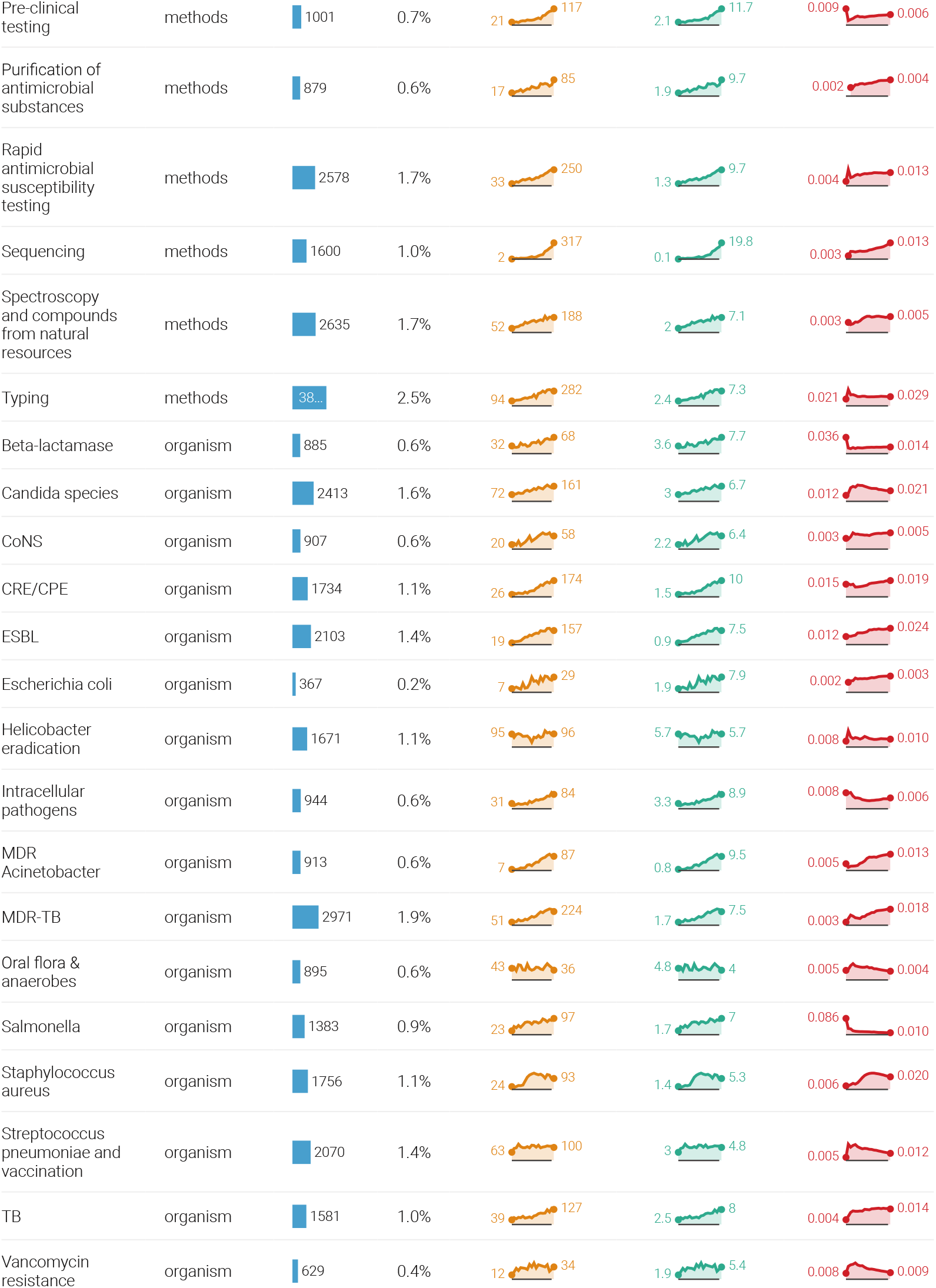
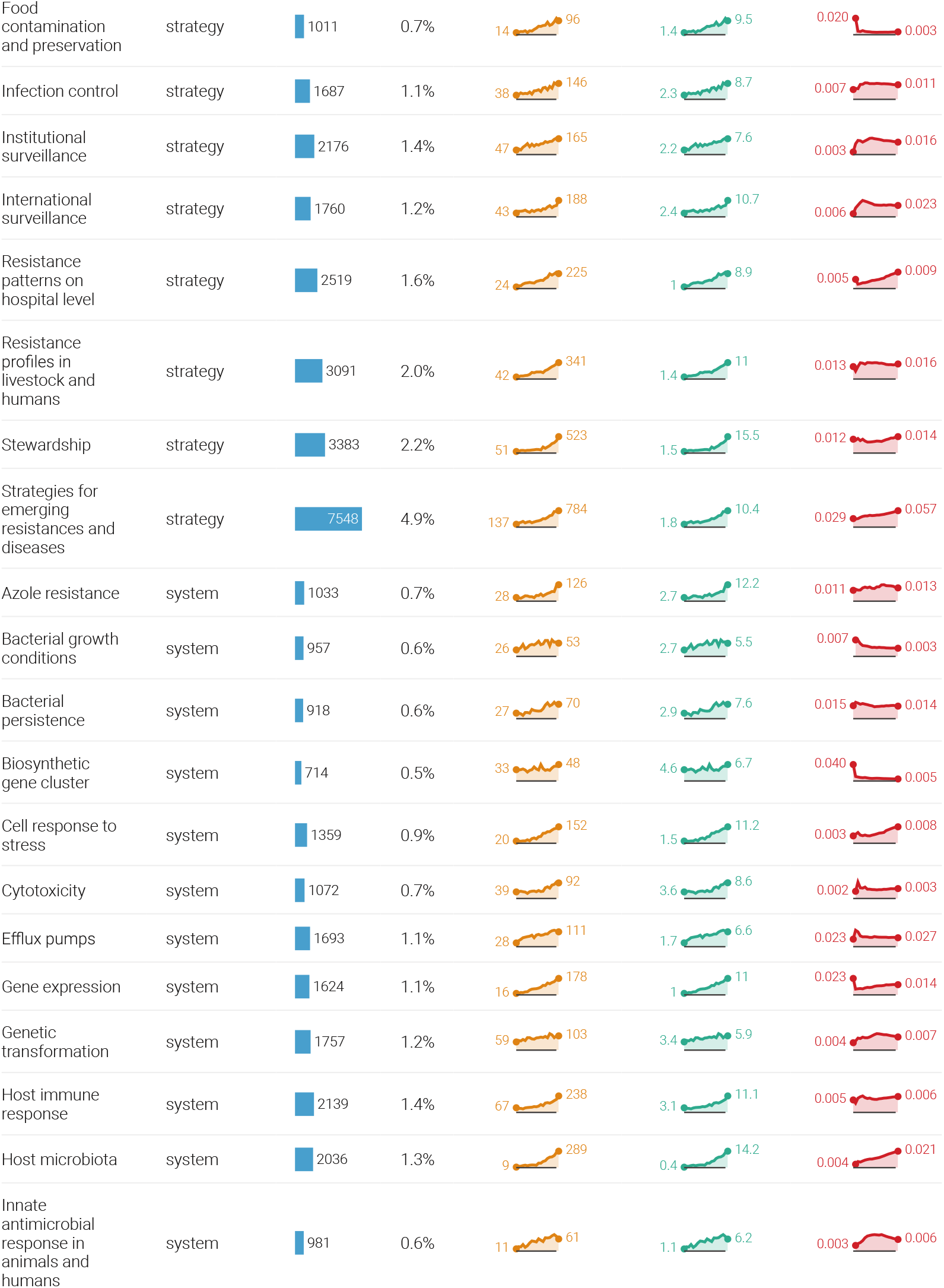
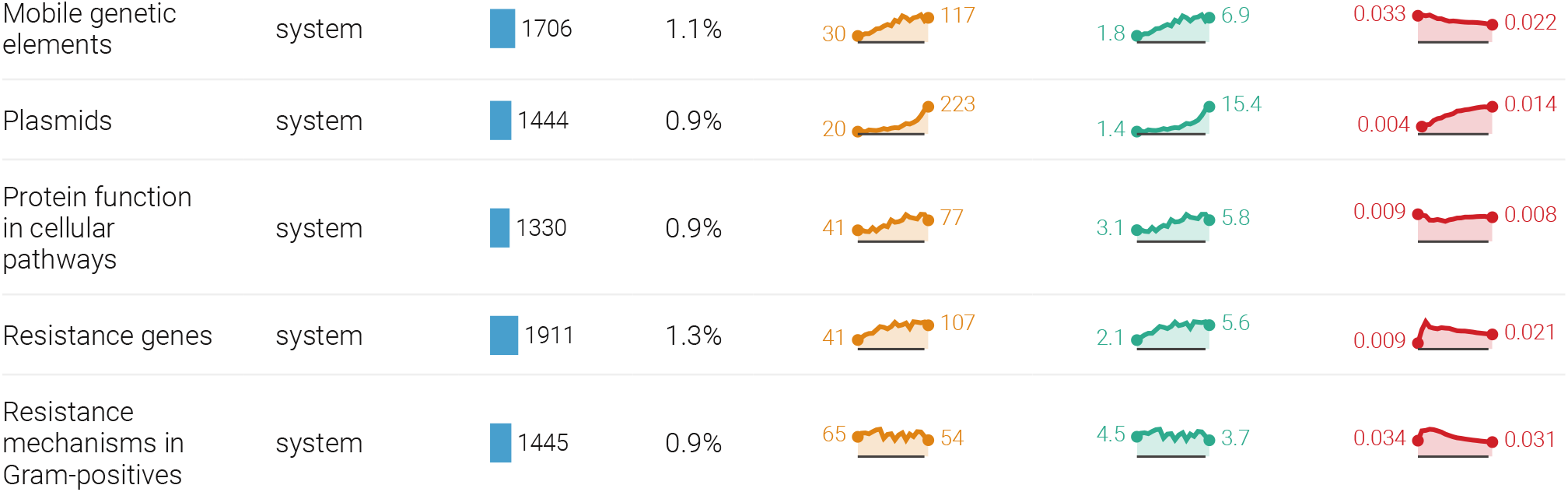
88 named topics in the field of AMR sorted alphabetically and by thematic group. Total number of publications per topic and in percent of all publications is presented. Trends in publication frequency, annual proportion, and PageRank are displayed for the entire period 1999-2018.

**Figure 4.**
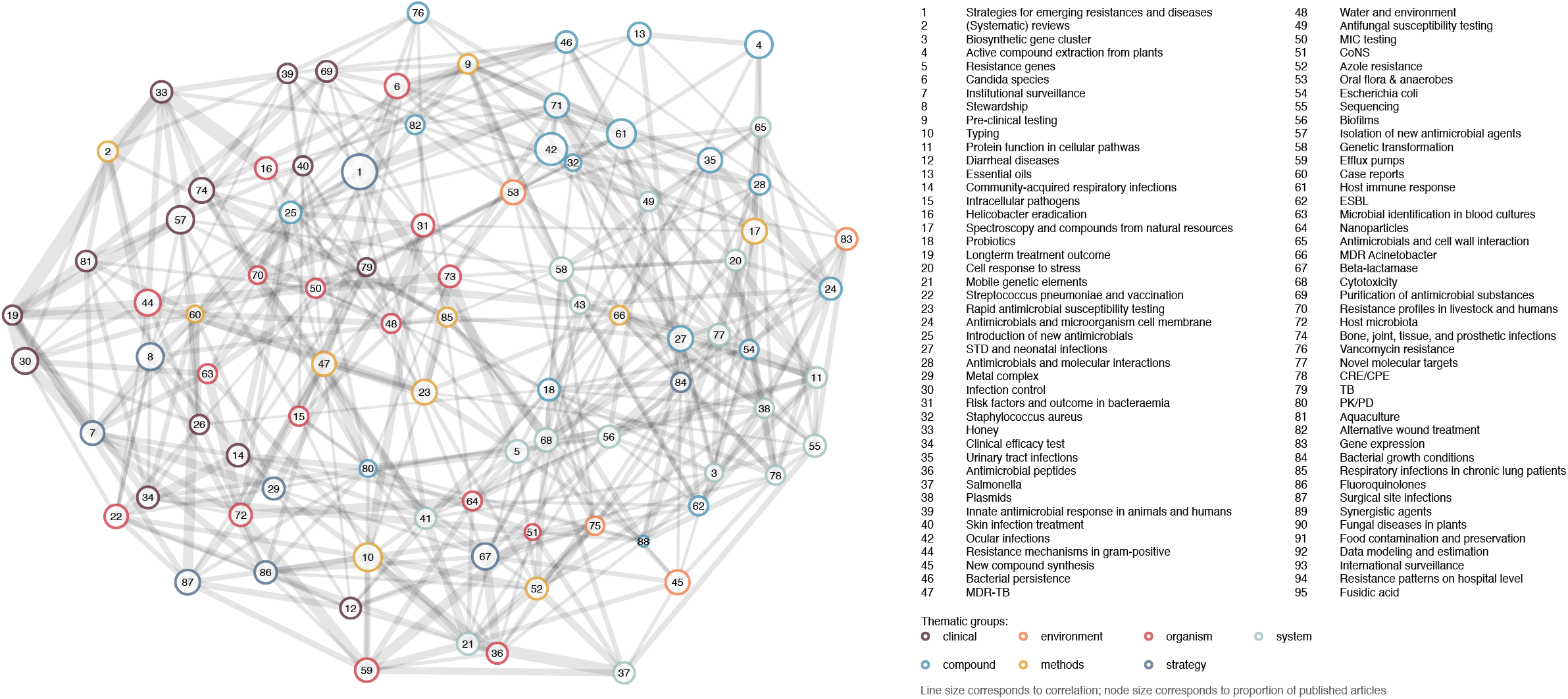
Topic network in AMR research based on topic co-occurrence generated. The 88 topics names with their corresponding number are displayed. Colours are used to highlight the 7 thematic groups within the network: clinical (purple), environment (orange), organism (red), system (grey-blue), compound (light blue), methods (yellow), and strategy (dark blue). Line size within the network corresponds to correlation weight (more weight equals thicker lines) and node size corresponds to the proportion of published articles.

## General trends

### Time

The absolute number and proportion of articles per topic underwent various changes over the last 20 years (Figure 2 and Figure 3). The most prevalent and important topics (top three), changes in prevalence and importance over the last 20 years, and publication bursts are highlighted in Figure 5.

**Figure 5.**
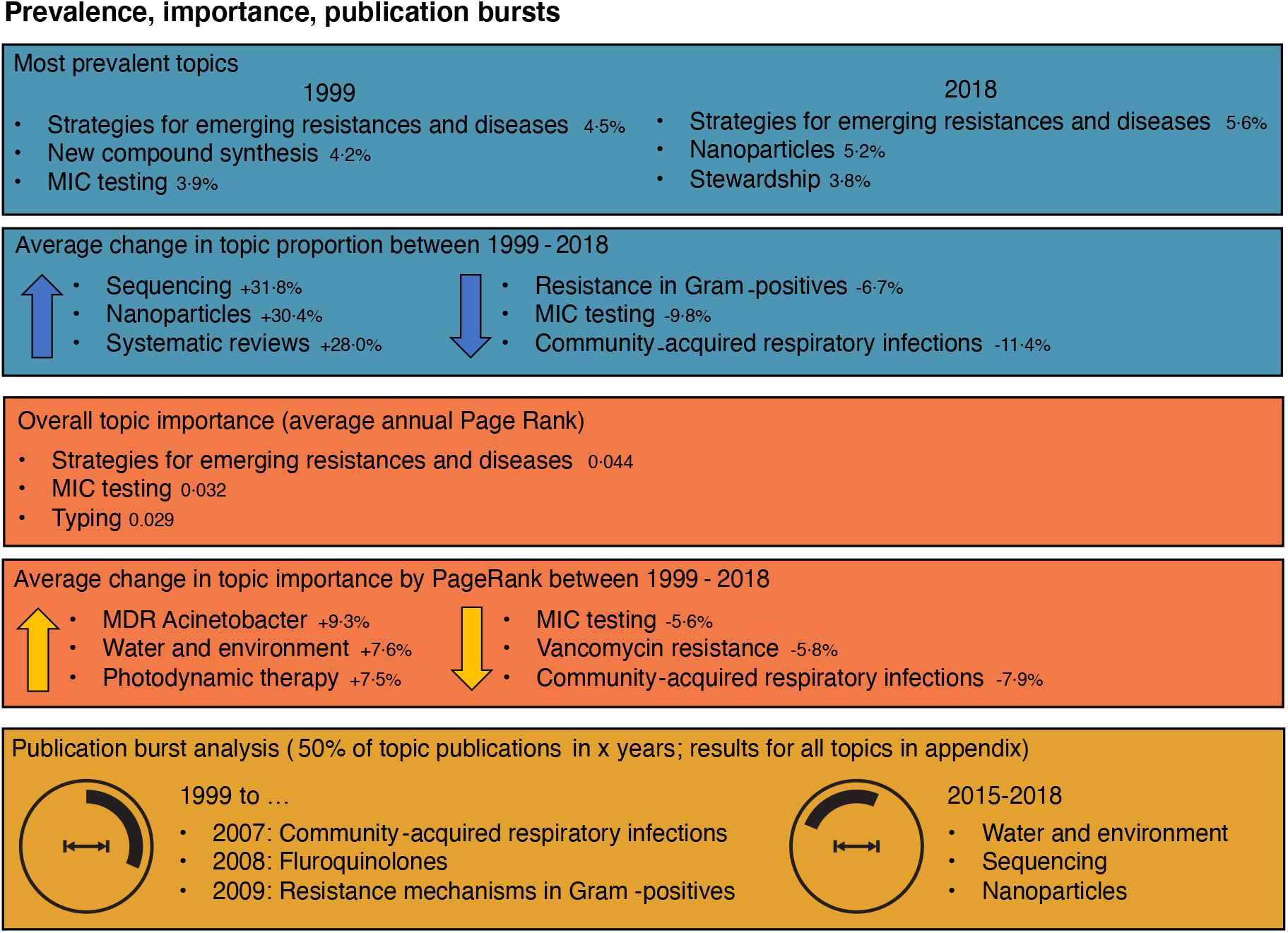
The most prevalent and important topics (top three) by: proportion of all published articles per year in 1999 and 2018 and average change (positive and negative) in proportion of all published articles between 1999 and 2018 (blue); average change (positive and negative) in topic importance (calculated PageRank) between 1999 and 2018 (orange); and publication bursts (period comprising 50% of all publications within one topic), longest and shortest intervals are shown (yellow).

### Geography

#### WHO regions

The first author’s country of affiliation could be extracted for 153 879 articles (97·0%) and 166 unique countries. Based on the country names, the results were stratified by WHO regions. The WHO regions of the Americas and the European Region were the largest overall contributors. However, other regions steadily increased their proportion of articles over time (Figure 6).

**Figure 6.**
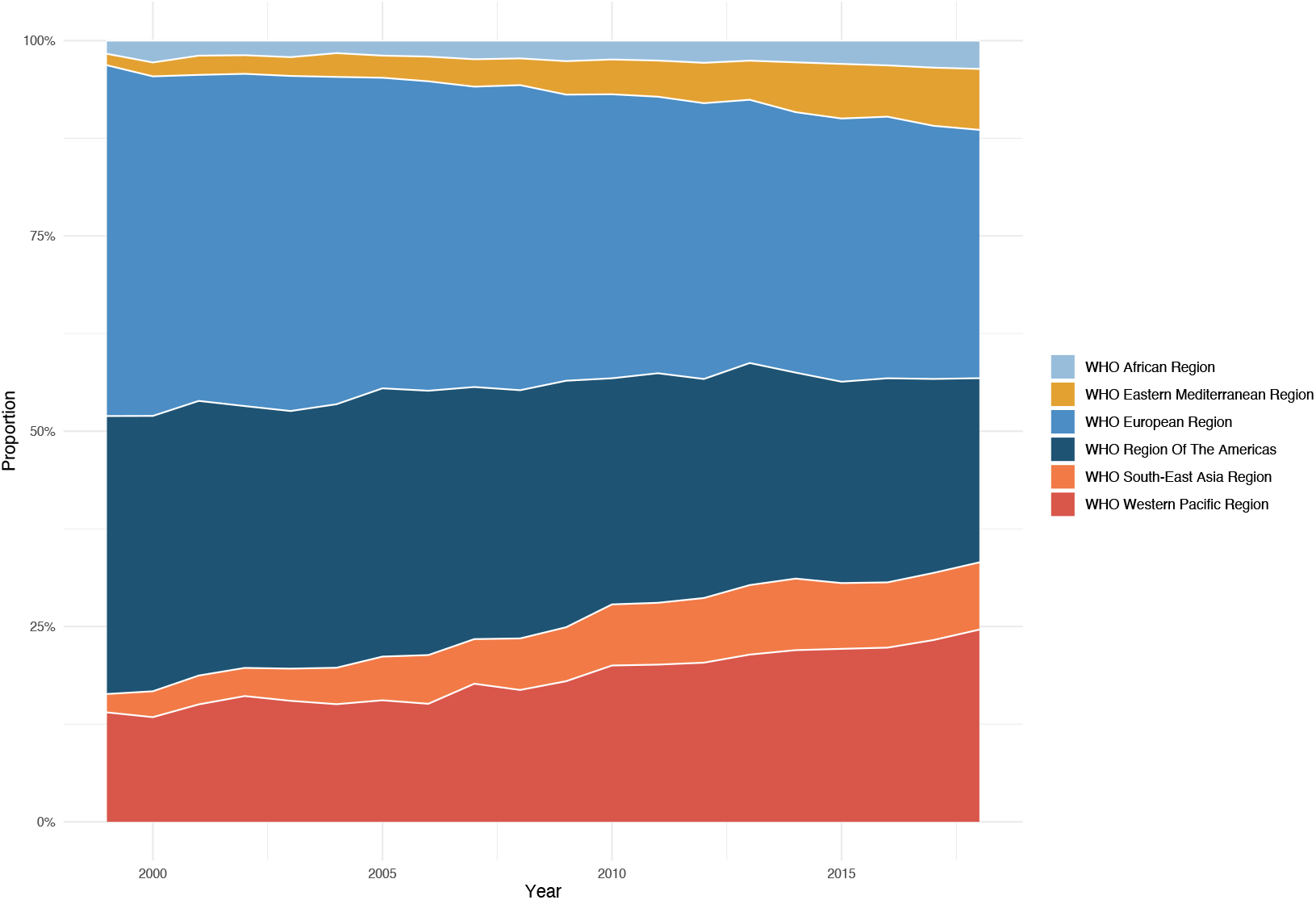
The annual proportion of AMR research per WHO region (1999-2018).

Varying patterns in the three most researched topics over time were observed per WHO region (Figure 7). While the European region and the region of the Americas focused strongly on *Strategies for emerging resistances and diseases*, the south-east Asian region and the western Pacific region demonstrated an increasing focus on *New compound synthesis* and *Nanoparticles. Resistance patterns on the hospital level* and *Active compound extraction from plants* were research priorities in the African and eastern Mediterranean region. Only the African region listed an organism-related topic (*MDR-TB*) among the top three topics. The importance of *MDR-TB* worldwide was further emphasized when assessing only organism-related topics per WHO region (Figure 8). All regions but the eastern Mediterranean listed *MDR-TB* in the top three of all organism-related topics over time.

**Figure 7.**
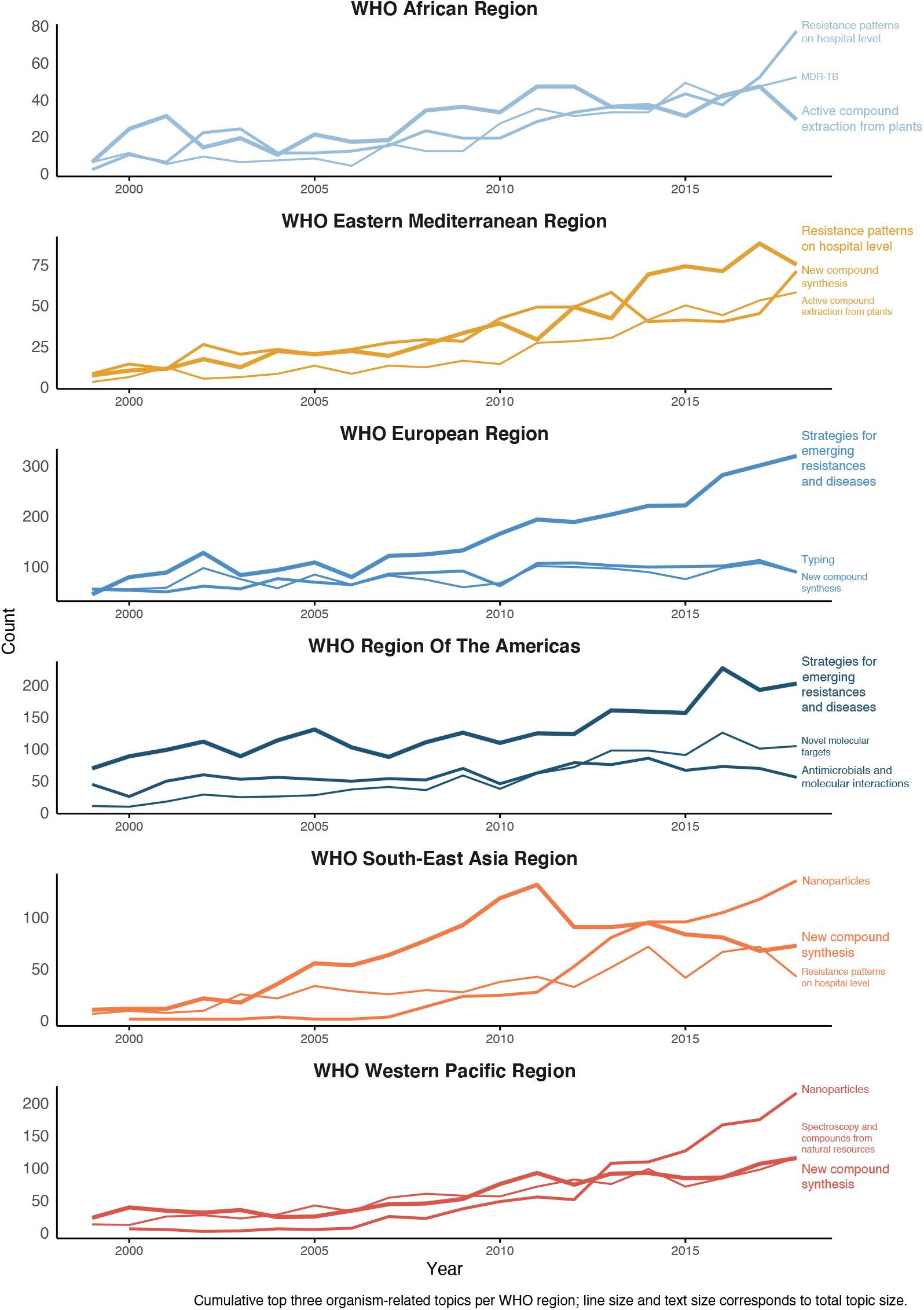
Research priorities within WHO regions per year (1999-2018). Top three topics by overall count per WHO region; line size and text size correspond to the total number of publications per topic and region.

**Figure 8.**
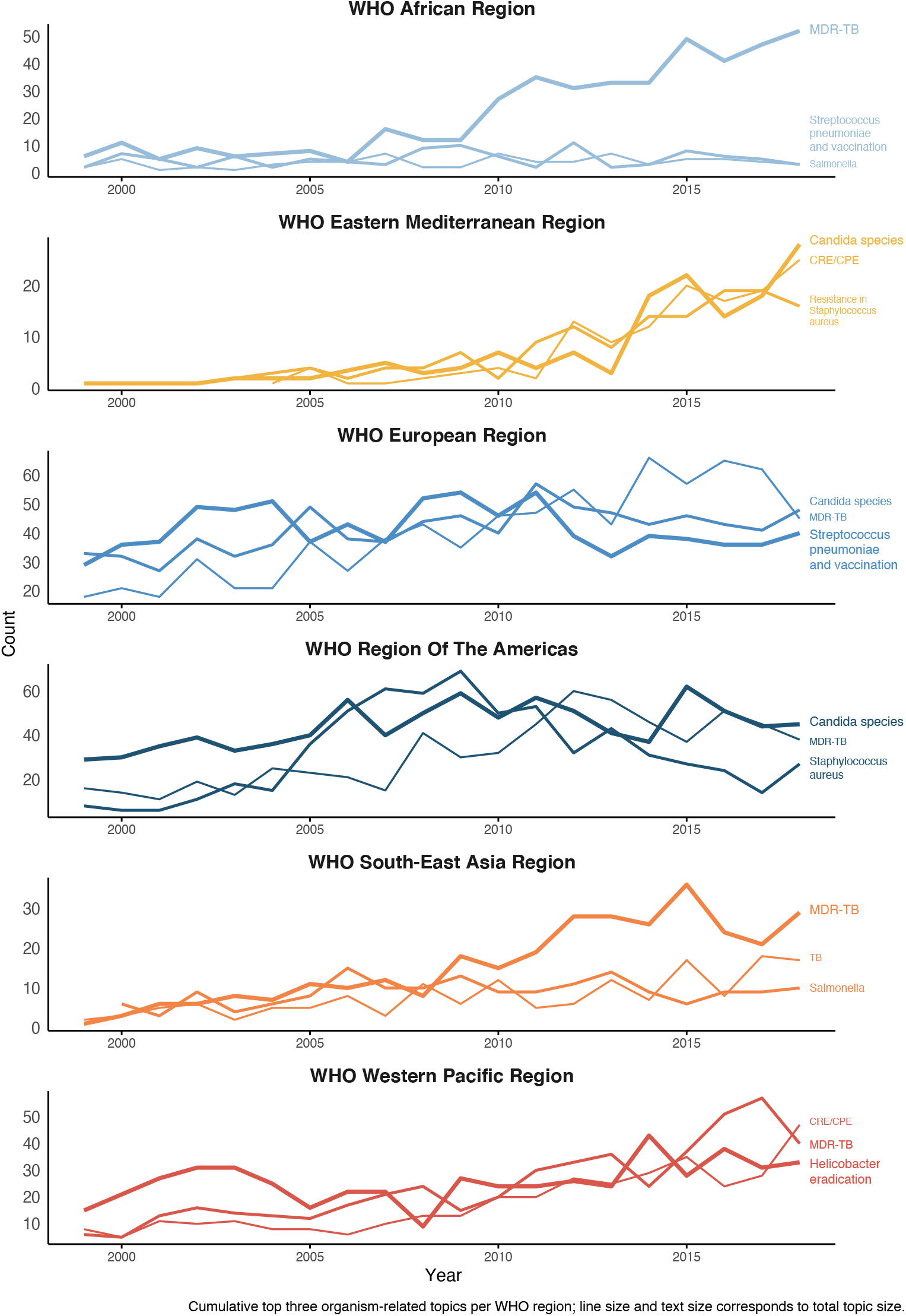
Organism-related research priorities within WHO regions per year. Top three organism-related topics by overall count per WHO region; line size and text size correspond to the total number of publications per topic and region.

#### Countries

At the country level, the United States of America (USA) contributed the most (ranked first in number of publications) to the body of AMR literature each year. However, China significantly increased its research output (n=2021 in 2018; +27·2% nominal annual increase), ranking second in 2018 after the USA (n=2092 in 2018; +4.9% nominal annual increase) and before India (n=1010 in 2018; +16·5% nominal annual increase). Regarding importance, according to PageRank per year, the USA led unchallenged throughout the years, followed by the United Kingdom (UK), France, Canada, Spain, and Germany (all countries in Appendix S6).

### Thematic groups

For the following sections, results were stratified into thematic groups based on topic names (Figure 3). Each thematic group will be introduced in terms of topic representation and elaborated on in terms of nominal and relative increase/decrease over time, importance based on PageRanks, and contributing countries.

#### Clinical-related theme

The clinical-related theme represents 14 topics related to clinical infections. Most clinical-related topics showed a steady increase over time, except for *Community-acquired respiratory infections*, which decreased after 2004. The most researched topic in 2018 was *Risk factors and outcome in bacteraemia* (n=266 articles), which steadily increased both nominally (+11·5% annually) and relatively (+3·0% annually) over the past 20 years (Figure 9). The topic importance by PageRank also increased over that period (+2·6% annually). In 2018, the countries who contributed most to *Risk factors and outcome in bacteraemia* were the USA (18·1%), China (14·6%), and the Republic of Korea (7·3%).

**Figure 9.**
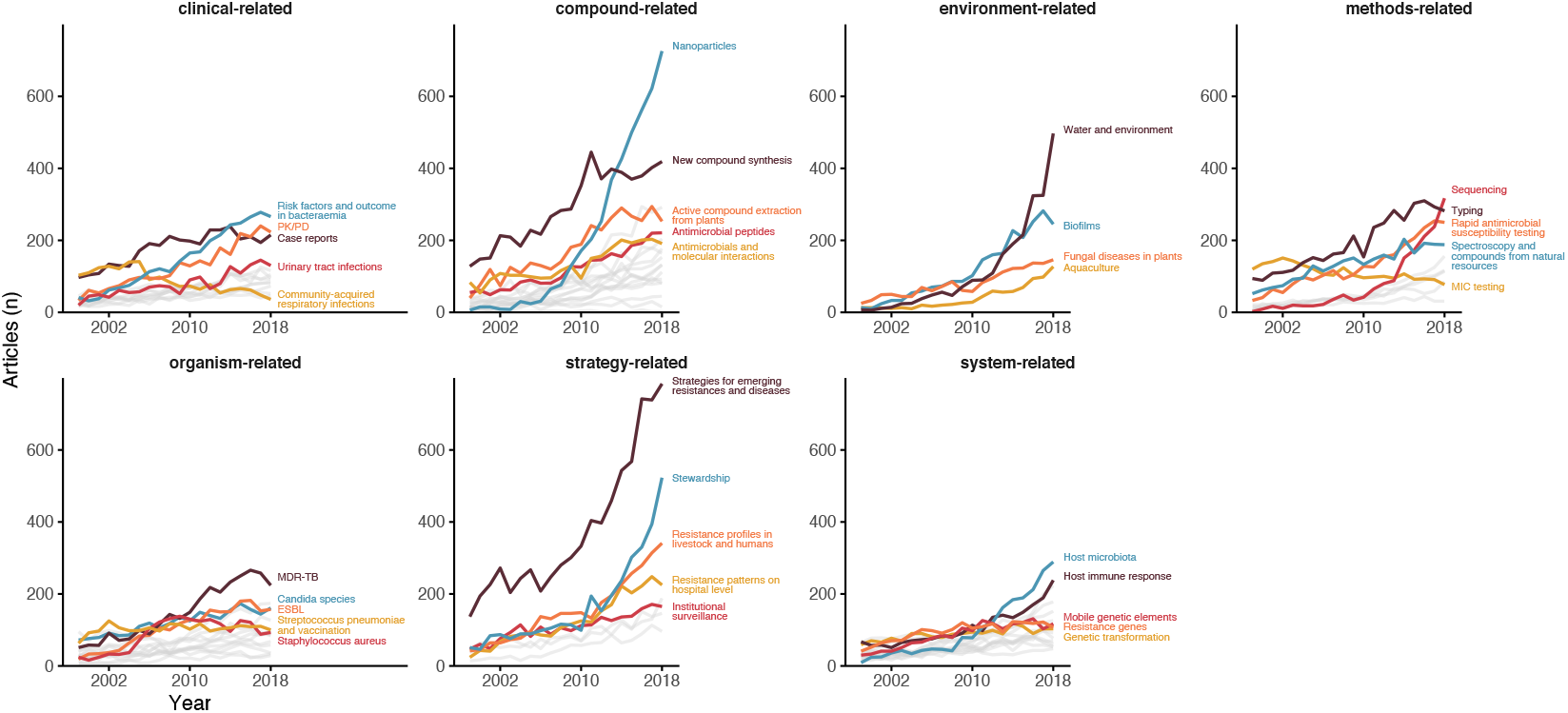
Five largest topics by total article count per thematic topic per year (1999-2018).

#### Compound-related theme

New antimicrobials and new compound strategies are grouped in the compound-related theme (18 topics). All compound-related topics showed a nominal increase over time, while the topic *Nanoparticles* showed a particularly steep increase after 2006 (32·6% mean annual increase) (Figure 9). In 2018, China and India were the largest contributors to *Nanoparticles*, ranking first (21·6%) and second (17·2%), respectively. A similar trend was observed for *New compound synthesis*. The largest contributors for *Active compound extraction from plants* were India (12·8%), Brazil (9·9%), and South Africa (5·6%). The USA was the largest contributor to *Antimicrobial peptides* and *Antimicrobials and molecular interactions*, although China ranked first in *Antimicrobial peptides* after 2015.

#### Environment-related theme

The environment-related theme comprises the topics *Water and environment, Biofilms, Aquaculture*, and *Fungal diseases in plants. Water and environment* showed a remarkable increase over time (+18·4% annual increase in proportion), ranking fourth in absolute and relative numbers in 2018 and also increased in importance by PageRank (+7·6% annual increase) (Figure 9). China mainly drove this trend with an almost exponential increase in the number of annual publications. In 2018, China contributed 40·9% of all publications on this topic. Moreover, *Water and environment* was China’s most researched topic after 2016. The topic was a driver for China’s increasing contribution to the overall body of AMR literature. The USA ranked second (12·1%) in *Water and environment* in 2018.

#### Methods-related theme

The methods-related theme represents 11 topics related to laboratory techniques and general research methodologies. *Typing* was the most prevalent method-related topic over 20 years (2·5% overall) but fell behind *Sequencing* in 2018, which showed a substantial increase after 2009 (+29·5% annually) (Figure 9). In the PageRank analysis, *Typing* ranked third in 2018 and demonstrated steady importance over the years. *Sequencing* ranked lower in the PageRank than *Typing* but gained importance over the years (+5·7% annual increase). In 2018, articles contributing to *Typing* came predominantly from China (20·9%), Brazil (7·8%), and the Republic of Korea (5·0%). In the same year, the USA (17·4%) was leading in *Sequencing* studies before China (13·6%) and the UK (6·0%).

#### Organism-related theme

Topics with a clear association to a specific organism or pathogen are grouped in the organism-related theme (16 topics). Of these topics, *MDR-TB* was most prevalent over time, with a peak in relative proportion in 2012 (10·8% of all topics). *Staphylococcus aureus* displayed a short but prominent peak in 2007-2008. The topics which increased the most in terms of the annual change in proportion were *Escherichia coli (+9·3%), MDR Acinetobacter (+9·0%), ESBL (+4·6%), CRE/CPE (+3·8%), and CoNS (+2·3%)*.

*MDR-TB* showed a distinct geographic distribution from the general trends in the AMR field. In 2018, most publications came from South Africa (11·6%), China (9·8%), and India (9·4%). Of the top ten countries with the highest burden of *MDR-TB*, according to the WHO, that were included in this study, 70% showed *MDR-TB* in their three most researched topics (Belarus, Kyrgyzstan, Republic of Moldova, Kazakhstan, Tajikistan, Uzbekistan, and Azerbaijan)^27^.

#### Strategy-related theme

The strategy-related theme represents topics related to conceptual strategies against AMR (eight topics). By far, the most prevalent topic was *Strategies for emerging resistances and diseases*, which also prevailed overall in AMR research (4·9% of total). The topic showed close links with all other topics as an overarching topic (see also citation cluster analysis in Appendix S4). *Stewardship, Institutional surveillance*, and *International surveillance* were the following three most prevalent strategy-related topics overall.

*Stewardship* was the third most researched topic in 2018 (3·8%), showing a remarkable increase over time (+16·1% annually) and in particular after 2010. Next to the USA, the UK significantly increased its contribution to *Stewardship*, ranking first in the topic in 2018 (16·8%). Within the UK, *Stewardship* was the most researched topic in 2018 (13·5%). Similar trends were identified for Australia and France, where *Stewardship* ranked first and second in 2018, respectively.

*International surveillance* and *Institutional surveillance* also increased in overall proportion but to a reduced extent (+1·7% and +0·5% annually, respectively). The importance by PageRank decreased by -2·4% annually for *International surveillance*, in contrast to an annual +1·3% increase for *Stewardship*. The USA (17·3%), the UK (10·9%), and Australia (7·3%) were the main contributors to *International surveillance. Institutional surveillance* showed similar trends to international surveillance but with a more dominant role for the USA (21·1%) as the primary contributor. Despite these similarities, the importance by PageRank for *Institutional surveillance* was consistently lower compared to *International surveillance*.

#### System-related theme

Molecular aspects and host characteristics are the focus of the 17 topics in the system-related theme. *Host microbiota* stood out with an annual nominal and relative increase of +24·0% and +14·5%, respectively (Figure 9). *Host microbiota* also gained importance over the years measured by PageRank (+6·3%). Other topics (*Mobile genetic elements, Efflux pumps, Resistance genes*, and *Genetic transformation*) did not show large variations in relative numbers over the years. The topics *Mobile genetic elements, Efflux pumps*, and *Resistance genes* were identified as important topics in terms of their PageRank over the years despite showing a steady decline. The USA was the largest contributor to most system-related topics, except for *Resistance genes* and *Mobile genetic elements*, where China was leading.

## Discussion

This study mapped 20 years (1999-2018) of AMR research using data-driven text-based techniques (structural topic modelling). We identified 88 topics across 166 countries. Topics, trends, and geographical differences were assessed. AMR publications increased by 450% over the two decades, and grew by 129% between 2004 and 2013 compared to 48.9% for all PubMed publications over the same period.^28^ The most prominent topics in 2018 were *Strategies for emerging resistances and diseases, Nanoparticles*, and *Stewardship*. Emerging topics included *Water and environment*, and *Sequencing*. Geographical trends highlighted the positive correlation between research on *MDR-TB* and the related MDR-TB burden.

### AMR research geography

The research geography changed remarkably over time, mainly due to increased contributions from India and China. The USA remained the leading country but showed a slower increase compared to other increasing countries. The geographical changes are similar to overall publication trends on PubMed, yet more pronounced.^28^ For example, China increased its overall research output by 271% on PubMed (2004-2013) compared to 609% for AMR over the same time.^28^

We identified research priorities on (supra-)national levels. Overall, the WHO European region and the region of the Americas produce much output on strategies and molecular aspects. *New compounds* and *Nanoparticles* were under focus in the South-East Asian and Western Pacific region. The African region is the only region listing an organism-specific topic, *MDR-TB*, in the top three researched topics. Considering organism-specific topics alone, *MDR-TB* played a significant role in most WHO regions and dominated organism-specific topics in the African region. It is positive to see that *MDR-TB* ranked high across AMR research output in the most affected countries. However, the overall trend for *MDR-TB* remained unchanged.

### General topic overview

We identified 88 topics that cover the diversity in AMR with high granularity. Previous more limited studies validate parts of the identified topics and our results overlap with a study that assessed topics in microbiology.^14^ Another study also assessed themes in microbiology and identified broad themes (e.g., animal models).^13^ Our results offer a greater granularity and a more extensive variety of topics (not exclusively microbiological), which can be grouped into these existing microbiological themes. Organism-specific trends were verified in the existing literature. *Staphylococcus aureus* showed an increase after 2004 with a peak in 2007/2008 and a decrease thereafter, as also observed by a scientometric study.^11^ Global surveillance data for methicillin-resistant *Staphylococcus aureus* (MRSA) also confirms this trend curve between the early 2000s and 2016.^29^ Among methods-related topics, *Sequencing* stood out and can be referred to as a “hot topic” based on publication bursts and PageRank, as confirmed by another topic modelling study on bioinformatics.^30^

### Limitations

Our study has several limitations. Foremost, topic modelling requires a manual selection of the number of topics (*K*). A higher value of *K* topics could have revealed additional topics. Furthermore, the risk of misclassified articles can not be fully eradicated as topic names were assigned based on the largest topic proportion identified per article. We extracted AMR publications using a holistic search definition but only one database was queried. The applied AMR definition comprised bacteria or fungi, whereas viruses and parasites were not included. Study inclusion was not limited by language as most indexed articles provide English titles and abstracts. Nevertheless, non-English publications might still be missing, potentially limiting geographical comparisons. Also, countries were determined by first author affiliation. This affiliation does not reflect the entire international network in the field. However, we hypothesize that the first authors might be closer to the geographic setting of the research focus.

### Future research

The generated data of this study enables detailed insights. Topic trends can be compared at the national level (Appendix S7), which was beyond the scope of this study. Moreover, leveraging additional data sources such as economic, funding, or diseases burden data could be used to study correlation effects. These future studies can deepen our understanding of the AMR field and streamline efforts to tackle current and future challenges. To this extent, all data are publicly available (https://osf.io/j3d65/) accompanied by an interactive analysis tool [topicsinamr.shinyapps.io/amr_topics]. We encourage readers to use the presented results to guide their analyses.

## Conclusion

We provide a comprehensive global map on important temporal and geographical trends in AMR over two decades. Using the entire AMR literature, an unprecedented data-driven approach identified several “hot” topics such as *Sequencing, Nanoparticles, Stewardship, Water and environment*, and *MDR-TB*. Simultaneously, the global research community has been changing and countries like China and India have become substantial contributors. This study and its publicly available data can be used to achieve a holistic view of the developments in the AMR field. Data on the global AMR burden is growing and information on AMR research funding is available. In the future, this can also be linked to data on global AMR research output, which has now been comprehensively assessed for the first time.

## Acknowledgements

We thank Jan Arends, Matthijs S. Berends, Francis F. Cavallo, Marjolein Heuker, and Nico Meessen for supporting the topic review and validation process.

