## Appendix for "Mapping Twenty Years of Antimicrobial Resistance Research Trends"

### S1. Stopwords

| | | | | | |
|---|---|---|---|---|---|
| resist | cent | includ | select | though | limit |
| antimicrobi | 2g | high | spp | even | altern |
| antimicrobialresist | per | mm | analysi | microgml | ms |
| antimicrob | can | allow | associ | gml | yearold |
| antibacterialresist | differ | display | ml | valu | reveal |
| antibacterialantifung | chang | exhibit | total | vs | compar |
| antibacteria | may | present | high | accord | pa |
| antibacteri | howev | found | low | perform | contain |
| antibiot | studi | show | result | trademark | cc |
| antibioticresist | trial | mgl | might | mgkg | ii |
| antifung | review | group | possibl | due | iii |
| drugresist | present | compare | prove | log | mgml |
| suscept | frequent | effect | demonstr | larg | shown |
| drugsuscept | two | known | confirm | suggest | conclud |
| bacteria | one | use | ci | systemat | appear |
| bacteri | three | spp | latter | search | suggest |
| 1 | higher | isol | investig | publish | thus |
| 2 | signific | standard | well | literatur | addit |
| 2mg | among | determin | correl | potenti | |
| 2m | mg | fit | first | avail | |

### S2. STM generative process

A probabilistic generative process is assumed to have generated the text in the corpus according to the process structure, associated parameters, and document-level covariates. Once the process is defined, the observed data are used to estimate the parameters used in the process using Bayesian inference.[39]

The probabilistic generative process is defined by letting each document in corpus with size $D$ be generated from $K$ distinct topics, consisting of a possible vocabulary of size $V$. Let $X$ be a $D \times P$ matrix containing document-level topic prevalence covariates as rows $x_d$. Similarly, let $Y$ be a $D \times A$ matrix containing document-level topical content variables as rows $y_d$.

Let each document $d$ have $N_d$ words indexed by $n$ such that $n \in \{1, ..., N_d\}$. Each $n$ is assigned a topic $z_{d,n}$ according to its assumed distribution

$$z_{d,n} | \theta_d \sim \text{Multinomial}(\theta_d)$$

Where $\theta_d$ is defined for each document $d$ as

$$\theta_d | X_d\gamma, \Sigma \sim \text{LogisticNormal}(\mu = X_d\gamma, \Sigma)$$

Where $\gamma$ is the $p \times 1$ coefficient vector and $\Sigma$ the $(K-1) \times (K-1)$ covariance matrix.

The probability distribution of the words in vocabulary $V$ for each topic $k$ and document $d$ with its associated covariates $X_d$ is given by

$$\beta_{d,k} \propto \exp(m + \kappa_k^{(t)} + \kappa_{y_d}^{(c)} + \kappa_{y_d,k}^{(i)}$$

where $m$ is the baseline word distribution and $\kappa_k^{(t)}$ and $\kappa_{y_d}^{(c)}$ are the topic-specific and covariate group deviations respectively and $\kappa_{y_d,k}^{(i)}$ is the interaction between the two.

For each $n$ word $w_{d,n}$ in document $d$ we sample a word from the multinomial distribution with parameter $\beta_{d,k}$ where $k = z_{d,n}$ defined as

$$w_{d,n} | z_{d,n}, \beta_{d,k=z_{d,n}} \sim \text{Multinomial}(\beta_{d,k=z_{d,n}})$$

The complete generative process is presented graphically in Fig. S3.

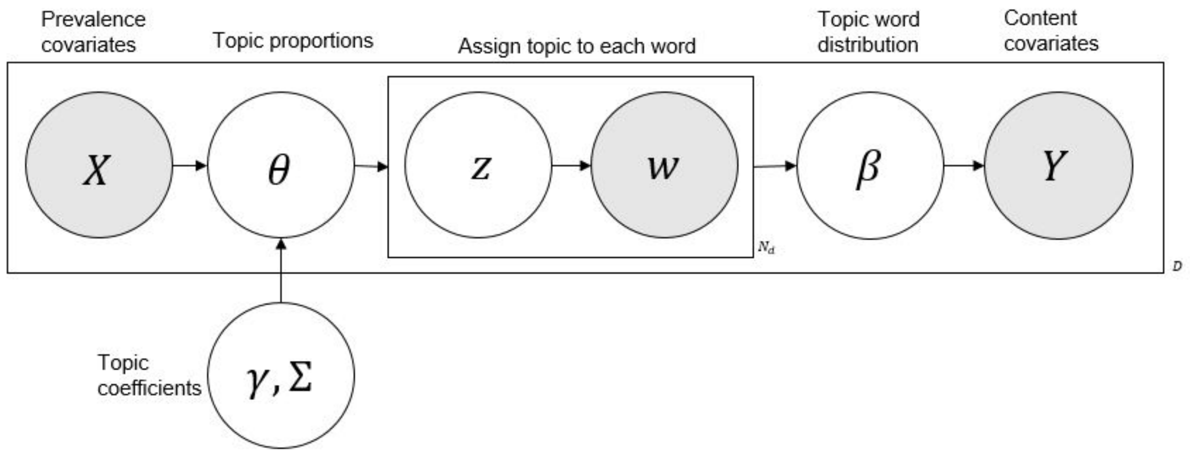


S3. A graphical illustration of the STM generative process with the observed variables shaded.

The parameters used in the generative process can be estimated using semi-collapsed variational expectation maximisation. The joint optimum for topic proportions $\theta_d$ and word-level topic assignment $z_{d,n}$ is calculated in the E-step while the global parameters ($\kappa, y$ and $\Sigma$) are inferred during the M-step.[39–42] The result is the estimated posterior distribution containing the multinomial distribution over the latent topics for each document and the multinomial distribution over the vocabulary for each of the identified topics.

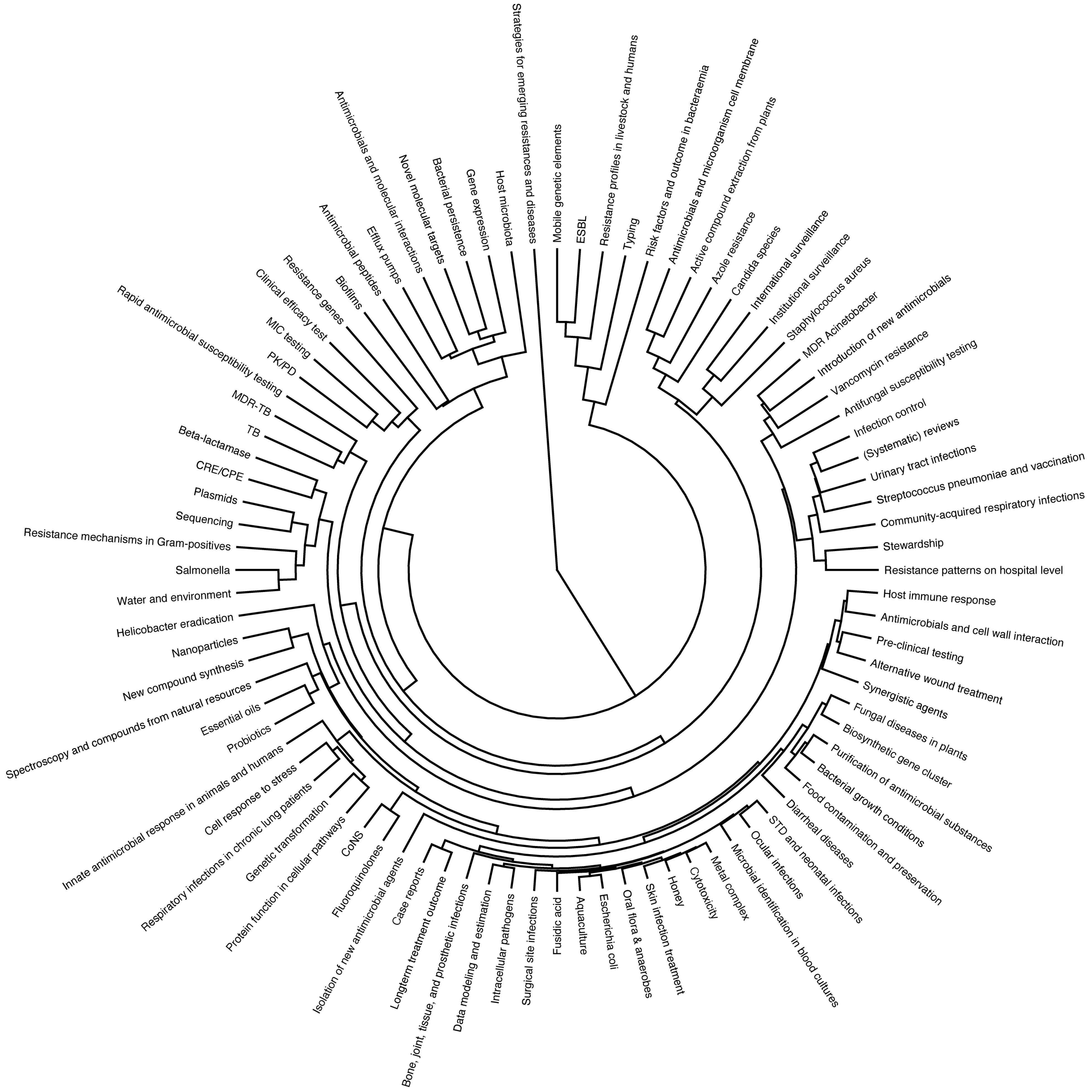


Figure S4. Topic correlation dendrogram based on topic citation data clustered using hierarchical clustering with Ward’s minimum variance method.

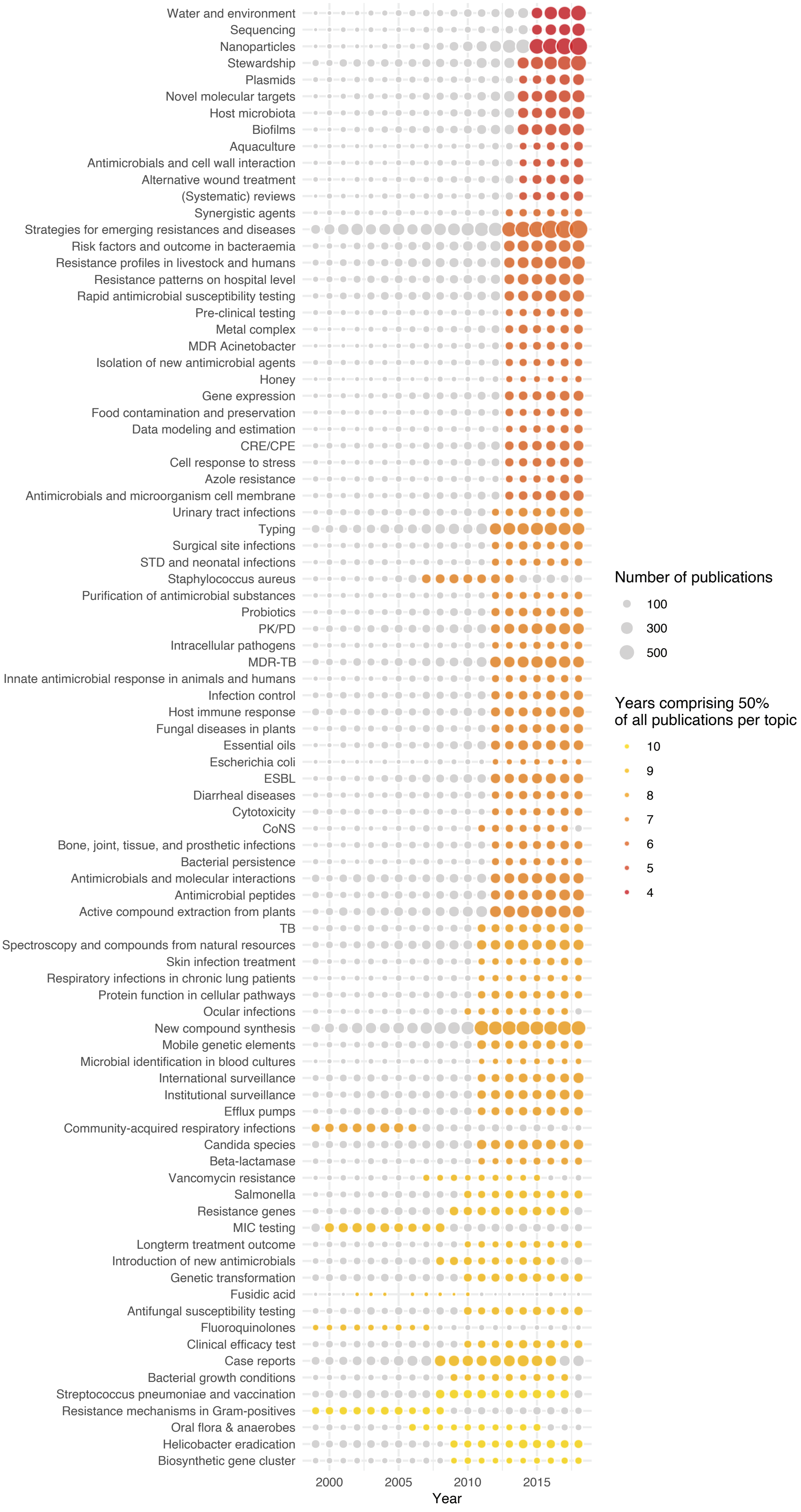

Water and environment
Sequencing
Nanoparticles
Stewardship
Plasmids
Novel molecular targets
Host microbiota
Biofilms
Aquaculture
Antimicrobials and cell wall interaction
Alternative wound treatment
(Systematic) reviews
Synergistic agents
Strategies for emerging resistances and diseases
Risk factors and outcome in bacteraemia
Resistance profiles in livestock and humans
Resistance patterns on hospital level
Rapid antimicrobial susceptibility testing
Pre-clinical testing
Metal complex
MDR Acinetobacter
Isolation of new antimicrobial agents
Honey
Gene expression
Food contamination and preservation
Data modeling and estimation
CRE/CPE
Cell response to stress
Azole resistance
Antimicrobials and microorganism cell membrane
Urinary tract infections
Typing
Surgical site infections
STD and neonatal infections
Staphylococcus aureus
Purification of antimicrobial substances
Probiotics
PK/PD
Intracellular pathogens
MDR-TB
Innate antimicrobial response in animals and humans
Infection control
Host immune response
Fungal diseases in plants
Essential oils
Escherichia coli
ESBL
Diarrheal diseases
Cytotoxicity
CoNS
Bone, joint, tissue, and prosthetic infections
Bacterial persistence
Antimicrobials and molecular interactions
Antimicrobial peptides
Active compound extraction from plants
TB
Spectroscopy and compounds from natural resources
Skin infection treatment
Respiratory infections in chronic lung patients
Protein function in cellular pathways
Ocular infections
New compound synthesis
Mobile genetic elements
Microbial identification in blood cultures
International surveillance
Institutional surveillance
Efflux pumps
Community-acquired respiratory infections
Candida species
Beta-lactamase
Vancomycin resistance
Salmonella
Resistance genes
MIC testing
Longterm treatment outcome
Introduction of new antimicrobials
Genetic transformation
Fusidic acid
Antifungal susceptibility testing
Fluoroquinolones
Clinical efficacy test
Case reports
Bacterial growth conditions
Streptococcus pneumoniae and vaccination
Resistance mechanisms in Gram-positives
Oral flora & anaerobes
Helicobacter eradication
Biosynthetic gene cluster
2000
2005
2010
2015
Year
Number of publications
100
300
500
Years comprising 50%
of all publications per topic
10
9
8
7
6
5
4

Figure S5. Publication burst, the least amount of years comprising 50% of all publications starting at the earliest year possible, calculated per topic. Bubble size represents the number of publications per year. Colours correspond to the number of years needed to comprise 50% of all publications per topic.

| Country code | WHO Region | n | % | Trend (n) | Trend (%) |
|---|---|---|---|---|---|
| AFG | WHO Eastern Mediterranean Region | 5 | 0,0% | 2 … 1 | 0,0% … 0,0% |
| AGO | WHO African Region | 1 | 0,0% | 1 | 0,0% |
| ALB | WHO European Region | 8 | 0,0% | 2 … 3 | 0,0% … 0,0% |
| ARE | WHO Eastern Mediterranean Region | 128 | 0,1% | 2 … 11 | 0,1% … 0,1% |
| ARG | WHO Region Of The Americas | 1055 | 0,7% | 15 … 89 | 0,5% … 0,6% |
| ARM | WHO European Region | 22 | 0,0% | 1 … 5 | 0,0% … 0,0% |
| ATG | WHO Region Of The Americas | 1 | 0,0% | 1 | 0,0% |
| AUS | WHO Western Pacific Region | 3093 | 2,0% | 46 … 354 | 1,5% … 2,5% |
| AUT | WHO European Region | 735 | 0,5% | 15 … 57 | 0,5% … 0,4% |
| AZE | WHO European Region | 5 | 0,0% | 1 … 2 | 0,0% … 0,0% |
| BDI | WHO African Region | 1 | 0,0% | 1 | 0,0% |
| BEL | WHO European Region | 1631 | 1,1% | 42 … 127 | 1,3% … 0,9% |
| BEN | WHO African Region | 34 | 0,0% | 3 … 3 | 0,0% … 0,0% |
| BFA | WHO African Region | 48 | 0,0% | 1 … 1 | 0,0% … 0,0% |
| BGD | WHO South-East Asia Region | 323 | 0,2% | 6 … 34 | 0,2% … 0,2% |
| BGR | WHO European Region | 192 | 0,1% | 8 … 11 | 0,3% … 0,1% |
| BHR | WHO Eastern Mediterranean Region | 17 | 0,0% | 1 … 1 | 0,0% … 0,0% |
| BIH | WHO European Region | 62 | 0,0% | 1 … 4 | 0,0% … 0,0% |
| BLR | WHO European Region | 19 | 0,0% | 1 … 1 | 0,0% … 0,0% |
| BOL | WHO Region Of The Americas | 4 | 0,0% | 1 … 1 | 0,0% … 0,0% |
| BRA | WHO Region Of The Americas | 4889 | 3,2% | 36 … 569 | 1,1% … 4,0% |
| BRB | WHO Region Of The Americas | 5 | 0,0% | 1 … 1 | 0,0% … 0,0% |
| BRN | WHO Western Pacific Region | 5 | 0,0% | 1 … 2 | 0,0% … 0,0% |
| BTN | WHO South-East Asia Region | 5 | 0,0% | 1 … 3 | 0,0% … 0,0% |
| BWA | WHO African Region | 36 | 0,0% | 2 … 2 | 0,1% … 0,0% |
| CAF | WHO African Region | 15 | 0,0% | 1 … 1 | 0,0% … 0,0% |

| Country code | WHO Region | n | % | Trend (n) | Trend (%) |
|---|---|---|---|---|---|
| CAN | WHO Region Of The Americas | 4221 | 2,7% | 131 336 | 4,2% 2,4% |
| CHE | WHO European Region | 1935 | 1,3% | 51 170 | 1,6% 1,2% |
| CHL | WHO Region Of The Americas | 398 | 0,3% | 5 49 | 0,2% 0,3% |
| CHN | WHO Western Pacific Region | 12139 | 7,9% | 32 2021 | 1,0% 14,2% |
| CIV | WHO African Region | 48 | 0,0% | 2 3 | 0,1% 0,0% |
| CMR | WHO African Region | 276 | 0,2% | 4 34 | 0,1% 0,2% |
| COD | WHO African Region | 26 | 0,0% | 1 2 | 0,0% 0,0% |
| COL | WHO Region Of The Americas | 413 | 0,3% | 3 40 | 0,1% 0,3% |
| CRI | WHO Region Of The Americas | 72 | 0,0% | 1 8 | 0,0% 0,1% |
| CUB | WHO Region Of The Americas | 101 | 0,1% | 5 3 | 0,2% 0,0% |
| CYP | WHO European Region | 19 | 0,0% | 1 2 | 0,0% 0,0% |
| CZE | WHO European Region | 733 | 0,5% | 6 59 | 0,2% 0,4% |
| DEU | WHO European Region | 5751 | 3,7% | 153 495 | 4,9% 3,5% |
| DJI | WHO Eastern Mediterranean Region | 1 | 0,0% | 1 | 0,0% |
| DMA | WHO Region Of The Americas | 1 | 0,0% | 1 | 0,0% |
| DNK | WHO European Region | 1357 | 0,9% | 40 112 | 1,3% 0,8% |
| DOM | WHO Region Of The Americas | 6 | 0,0% | 3 1 | 0,0% 0,0% |
| DZA | WHO African Region | 249 | 0,2% | 1 38 | 0,0% 0,3% |
| ECU | WHO Region Of The Americas | 43 | 0,0% | 1 16 | 0,0% 0,1% |
| EGY | WHO Eastern Mediterranean Region | 1242 | 0,8% | 12 181 | 0,4% 1,3% |
| ERI | WHO African Region | 6 | 0,0% | 1 2 | 0,0% 0,0% |
| ESP | WHO European Region | 5623 | 3,7% | 140 401 | 4,5% 2,8% |
| EST | WHO European Region | 91 | 0,1% | 1 9 | 0,0% 0,1% |
| ETH | WHO African Region | 403 | 0,3% | 4 89 | 0,1% 0,6% |
| FIN | WHO European Region | 545 | 0,4% | 24 37 | 0,8% 0,3% |
| FJI | WHO Western Pacific Region | 9 | 0,0% | 1 1 | 0,0% 0,0% |
| FRA | WHO European Region | 5793 | 3,8% | 215 412 | 6,8% 2,9% |

| Country code | WHO Region | n | % | Trend (n) | Trend (%) |
|---|---|---|---|---|---|
| GAB | WHO African Region | 19 | 0,0% | 1 6 | 0,0% 0,1% |
| GBR | WHO European Region | 7394 | 4,8% | 215 652 | 6,8% 4,6% |
| GEO | WHO European Region | 814 | 0,5% | 24 92 | 0,8% 0,6% |
| GHA | WHO African Region | 115 | 0,1% | 1 21 | 0,0% 0,1% |
| GIN | WHO African Region | 27 | 0,0% | 12 2 | 0,3% 0,0% |
| GMB | WHO African Region | 15 | 0,0% | 1 1 | 0,0% 0,0% |
| GNB | WHO African Region | 1 | 0,0% | 1 | 0,0% |
| GRC | WHO European Region | 1305 | 0,8% | 15 77 | 0,5% 0,5% |
| GRD | WHO Region Of The Americas | 17 | 0,0% | 1 2 | 0,0% 0,0% |
| GTM | WHO Region Of The Americas | 9 | 0,0% | 1 2 | 0,0% 0,0% |
| GUY | WHO Region Of The Americas | 3 | 0,0% | 1 1 | 0,0% 0,0% |
| HND | WHO Region Of The Americas | 5 | 0,0% | 1 2 | 0,0% 0,0% |
| HRV | WHO European Region | 307 | 0,2% | 3 31 | 0,1% 0,2% |
| HTI | WHO Region Of The Americas | 8 | 0,0% | 1 2 | 0,0% 0,0% |
| HUN | WHO European Region | 503 | 0,3% | 8 45 | 0,3% 0,3% |
| IDN | WHO South-East Asia Region | 127 | 0,1% | 1 24 | 0,0% 0,2% |
| IND | WHO South-East Asia Region | 9195 | 6,0% | 60 1010 | 1,9% 7,1% |
| IRL | WHO European Region | 856 | 0,6% | 11 67 | 0,4% 0,5% |
| IRN | WHO Eastern Mediterranean Region | 2619 | 1,7% | 2 405 | 0,1% 2,8% |
| IRQ | WHO Eastern Mediterranean Region | 86 | 0,1% | 2 16 | 0,0% 0,1% |
| ISL | WHO European Region | 162 | 0,1% | 4 26 | 0,1% 0,2% |
| ISR | WHO European Region | 1298 | 0,8% | 30 96 | 1,0% 0,7% |
| ITA | WHO European Region | 5320 | 3,5% | 127 483 | 4,0% 3,4% |
| JAM | WHO Region Of The Americas | 67 | 0,0% | 1 4 | 0,0% 0,0% |
| JOR | WHO Eastern Mediterranean Region | 206 | 0,1% | 3 24 | 0,1% 0,2% |
| JPN | WHO Western Pacific Region | 6484 | 4,2% | 239 369 | 7,6% 2,6% |

| Country code | WHO Region | n | % | Trend (n) | Trend (%) |
|---|---|---|---|---|---|
| KAZ | WHO European Region | 29 | 0,0% | 1 6 | 0,0% 0,0% |
| KEN | WHO African Region | 187 | 0,1% | 2 15 | 0,1% 0,1% |
| KGZ | WHO European Region | 4 | 0,0% | 1 1 | 0,0% 0,0% |
| KHM | WHO Western Pacific Region | 29 | 0,0% | 1 1 | 0,0% 0,0% |
| KNA | WHO Region Of The Americas | 2 | 0,0% | 1 1 | 0,0% 0,0% |
| KOR | WHO Western Pacific Region | 3875 | 2,5% | 52 369 | 1,7% 2,6% |
| KWT | WHO Eastern Mediterranean Region | 231 | 0,2% | 5 14 | 0,2% 0,1% |
| LAO | WHO Western Pacific Region | 12 | 0,0% | 1 1 | 0,0% 0,0% |
| LBN | WHO Eastern Mediterranean Region | 251 | 0,2% | 4 44 | 0,1% 0,3% |
| LBR | WHO African Region | 1 | 0,0% | 1 | 0,0% |
| LBY | WHO Eastern Mediterranean Region | 48 | 0,0% | 2 2 | 0,1% 0,0% |
| LKA | WHO South-East Asia Region | 76 | 0,0% | 2 15 | 0,1% 0,1% |
| LSO | WHO African Region | 4 | 0,0% | 3 1 | 0,0% 0,0% |
| LTU | WHO European Region | 135 | 0,1% | 2 14 | 0,1% 0,1% |
| LUX | WHO European Region | 15 | 0,0% | 1 1 | 0,0% 0,0% |
| LVA | WHO European Region | 34 | 0,0% | 1 9 | 0,0% 0,1% |
| MAR | WHO Eastern Mediterranean Region | 218 | 0,1% | 2 24 | 0,1% 0,2% |
| MDA | WHO European Region | 6 | 0,0% | 1 1 | 0,0% 0,0% |
| MDG | WHO African Region | 34 | 0,0% | 1 4 | 0,0% 0,0% |
| MEX | WHO Region Of The Americas | 1033 | 0,7% | 13 114 | 0,4% 0,8% |
| MKD | WHO European Region | 11 | 0,0% | 1 1 | 0,0% 0,0% |
| MLI | WHO African Region | 15 | 0,0% | 1 2 | 0,0% 0,0% |
| MLT | WHO European Region | 25 | 0,0% | 1 2 | 0,0% 0,0% |
| MMR | WHO South-East Asia Region | 16 | 0,0% | 1 5 | 0,0% 0,0% |
| MNE | WHO European Region | 4 | 0,0% | 1 1 | 0,0% 0,0% |
| MNG | WHO Western Pacific Region | 30 | 0,0% | 1 2 | 0,0% 0,0% |

| Country code | WHO Region | n | % | Trend (n) | Trend (%) |
|---|---|---|---|---|---|
| MOZ | WHO African Region | 26 | 0,0% | 1 6 | 0,0% 0,0% |
| MRT | WHO African Region | 2 | 0,0% | 1 1 | 0,0% 0,0% |
| MUS | WHO African Region | 24 | 0,0% | 1 2 | 0,0% 0,0% |
| MWI | WHO African Region | 46 | 0,0% | 1 3 | 0,0% 0,0% |
| MYS | WHO Western Pacific Region | 871 | 0,6% | 7 85 | 0,2% 0,6% |
| NAM | WHO African Region | 14 | 0,0% | 1 1 | 0,0% 0,0% |
| NER | WHO African Region | 9 | 0,0% | 1 2 | 0,0% 0,0% |
| NGA | WHO African Region | 635 | 0,4% | 10 69 | 0,3% 0,5% |
| NIC | WHO Region Of The Americas | 6 | 0,0% | 2 1 | 0,1% 0,0% |
| NLD | WHO European Region | 1988 | 1,3% | 77 124 | 2,5% 0,9% |
| NOR | WHO European Region | 684 | 0,4% | 24 41 | 0,8% 0,3% |
| NPL | WHO South-East Asia Region | 234 | 0,2% | 2 22 | 0,1% 0,2% |
| NZL | WHO Western Pacific Region | 496 | 0,3% | 17 35 | 0,5% 0,2% |
| OMN | WHO Eastern Mediterranean Region | 59 | 0,0% | 2 2 | 0,1% 0,0% |
| PAK | WHO Eastern Mediterranean Region | 1112 | 0,7% | 8 193 | 0,3% 1,4% |
| PAN | WHO Region Of The Americas | 16 | 0,0% | 1 2 | 0,0% 0,0% |
| PER | WHO Region Of The Americas | 145 | 0,1% | 2 19 | 0,1% 0,1% |
| PHL | WHO Western Pacific Region | 80 | 0,1% | 1 2 | 0,0% 0,0% |
| PNG | WHO Western Pacific Region | 14 | 0,0% | 1 4 | 0,0% 0,0% |
| POL | WHO European Region | 2191 | 1,4% | 26 213 | 0,8% 1,5% |
| PRK | WHO South-East Asia Region | 1 | 0,0% | 1 | 0,0% |
| PRT | WHO European Region | 1561 | 1,0% | 18 160 | 0,6% 1,1% |
| PRY | WHO Region Of The Americas | 17 | 0,0% | 1 2 | 0,0% 0,0% |
| PSE | WHO Eastern Mediterranean Region | 57 | 0,0% | 1 3 | 0,0% 0,0% |
| QAT | WHO Eastern Mediterranean Region | 53 | 0,0% | 3 8 | 0,1% 0,1% |

| Country code | WHO Region | n | % | Trend (n) | Trend (%) |
|---|---|---|---|---|---|
| ROU | WHO European Region | 461 | 0,3% | 3 45 | 0,1% 0,3% |
| RUS | WHO European Region | 723 | 0,5% | 27 77 | 0,9% 0,5% |
| RWA | WHO African Region | 9 | 0,0% | 3 2 | 0,0% 0,0% |
| SAU | WHO Eastern Mediterranean Region | 749 | 0,5% | 5 109 | 0,2% 0,8% |
| SDN | WHO Eastern Mediterranean Region | 50 | 0,0% | 1 7 | 0,0% 0,0% |
| SEN | WHO African Region | 65 | 0,0% | 7 2 | 0,2% 0,0% |
| SGP | WHO Western Pacific Region | 714 | 0,5% | 3 79 | 0,1% 0,6% |
| SLV | WHO Region Of The Americas | 2 | 0,0% | 1 1 | 0,0% 0,0% |
| SRB | WHO European Region | 365 | 0,2% | 4 39 | 0,1% 0,3% |
| SVK | WHO European Region | 324 | 0,2% | 19 13 | 0,6% 0,1% |
| SVN | WHO European Region | 218 | 0,1% | 2 21 | 0,1% 0,1% |
| SWE | WHO European Region | 1654 | 1,1% | 47 130 | 1,5% 0,9% |
| SWZ | WHO African Region | 2 | 0,0% | 1 1 | 0,0% 0,0% |
| SYR | WHO Eastern Mediterranean Region | 17 | 0,0% | 1 1 | 0,0% 0,0% |
| TCD | WHO African Region | 7 | 0,0% | 1 1 | 0,0% 0,0% |
| TGO | WHO African Region | 13 | 0,0% | 1 2 | 0,0% 0,0% |
| THA | WHO South-East Asia Region | 1173 | 0,8% | 5 115 | 0,2% 0,8% |
| TJK | WHO European Region | 1 | 0,0% | 1 | 0,0% |
| TTO | WHO Region Of The Americas | 53 | 0,0% | 3 4 | 0,1% 0,0% |
| TUN | WHO Eastern Mediterranean Region | 590 | 0,4% | 3 65 | 0,1% 0,5% |
| TUR | WHO European Region | 2646 | 1,7% | 24 151 | 0,8% 1,1% |
| TWN | WHO Western Pacific Region | 2394 | 1,6% | 40 158 | 1,3% 1,1% |
| TZA | WHO African Region | 142 | 0,1% | 3 23 | 0,1% 0,2% |
| UGA | WHO African Region | 119 | 0,1% | 3 14 | 0,1% 0,1% |
| UKR | WHO European Region | 110 | 0,1% | 5 12 | 0,2% 0,1% |

| Country code | WHO Region | n | % | Trend (n) | Trend (%) |
|---|---|---|---|---|---|
| URY | WHO Region Of The Americas | 80 | 0,1% | 1 6 | 0,0% 0,0% |
| USA | WHO Region Of The Americas | 31871 | 20,7% | 901 2092 | 28,7% 14,7% |
| UZB | WHO European Region | 10 | 0,0% | 1 2 | 0,0% 0,0% |
| VEN | WHO Region Of The Americas | 126 | 0,1% | 2 6 | 0,1% 0,0% |
| VNM | WHO Western Pacific Region | 223 | 0,1% | 4 27 | 0,1% 0,2% |
| XKX | WHO European Region | 12 | 0,0% | 1 2 | 0,0% 0,0% |
| YEM | WHO Eastern Mediterranean Region | 30 | 0,0% | 1 3 | 0,0% 0,0% |
| ZAF | WHO African Region | 1393 | 0,9% | 28 170 | 0,9% 1,2% |
| ZMB | WHO African Region | 27 | 0,0% | 2 1 | 0,0% 0,0% |
| ZWE | WHO African Region | 38 | 0,0% | 2 3 | 0,1% 0,0% |

Figure S6. 166 contributing countries sorted alphabetically by country code. Total number of publications per country and in percent of all publications is presented. Trends in publication frequency and annual proportion are displayed for the entire period 1999-2018.

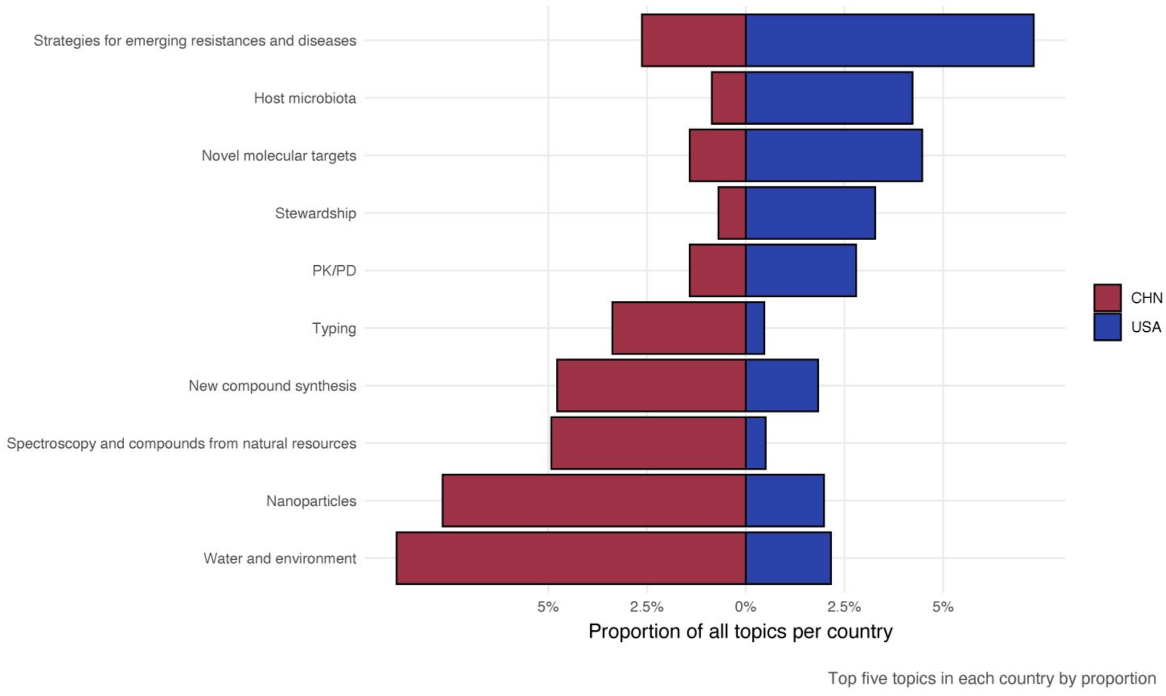


Figure S7. Topic distribution of top five topics within China and the USA (2016-2018).
